# Phenome-wide association study of a comprehensive health check-up database in a Korea population: Clinical application & trans-ethnic comparison

**DOI:** 10.1101/2020.05.31.126201

**Authors:** Eun Kyung Choe, Manu Shivakumar, Anurag Verma, Shefali Setia Verma, Seung Ho Choi, Joo Sung Kim, Dokyoon Kim

## Abstract

**Background:** The expanding use of the phenome-wide association study (PheWAS) faces challenges in the context of using International Classification of Diseases billing codes for phenotype definition, imbalanced study population ethnicity, and constrained application of the results to clinical practice or research.

**Methods:** We performed a PheWAS utilizing deep phenotypes corroborated by comprehensive health check-ups in a Korean population, along with trans-ethnic comparisons through the UK Biobank and Biobank Japan Project. Network analysis, visualization of cross-phenotype mapping, and causal inference mapping with Mendelian randomization were conducted in order to make robust, clinically applicable interpretations.

**Results:** Of the 136 phenotypes extracted from the health check-up database, the PheWAS associated 65 phenotypes with 14,101 significant variants (*P* < 4.92×10^−10^). In the association study for body mass index, our population showed 583 exclusive loci relative to the Japanese population and 669 exclusive loci relative to the European population. In the meta-analysis with Korean and Japanese populations, 72.5% of phenotypes had uniquely significant variants. Tumor markers and hematologic phenotypes had a high degree of phenotype-phenotype pairs. By Mendelian randomization, one skeletal muscle mass phenotype was causal and two were outcomes. Among phenotype pairs from the genotype-driven cross-phenotype associations, 71.65% also demonstrated penetrance in correlation analysis using a clinical database.

**Conclusions:** This comprehensive analysis of PheWAS results based on a health check-up database will provide researchers and clinicians with a panoramic overview of the networks among multiple phenotypes and genetic variants, laying groundwork for the practical application of precision medicine.

## Introduction

From the healthcare perspective, the key concept of precision medicine generally refers to incorporating genetic, lifestyle, environmental, and cultural factors into one’s health status for the purpose of promoting health, improving diagnostic accuracy, preventing disease, and providing personalized treatment ^1,2^. To make precision medicine functional, a systematic and integrational understanding of each individual’s clinical and genetic information is required. The phenome-wide association study (PheWAS) is one tool able to fulfill the purpose of precision medicine ^3^. PheWAS explores associations among genetic variants and a wide range of traits, including clinical outcomes and lifestyle, environmental, and cultural factors ^4^. The rapid growth of genomic resources linked to electronic health record (EHR) databases has increasingly become leveraged by PheWAS ^5^. In laying the foundation for precision medicine, PheWAS has several advantages. It could identify pleiotropic effects (cross-phenotype associations) ^6^ and shared genetic factors that influence comorbidities or influential factors associated with phenotypes ^3,7^ or laboratory results ^8,9^, which can eventually enable a personalized healthcare approach ^7^.

However, PheWAS to date has encountered several challenges in practice. First, most PheWASs defined phenotypes using International Classification of Diseases (ICD) terms such as billing codes or phecodes (a type of ICD code grouping). These billing codes can bring an underlying bias into healthcare practices ^5,10^. Second, most genetic association studies have been done in limited, non-Asian populations ^10^. A PheWAS performed on a homogeneous population from a singular nation can be more powerful as the pools of cases and controls are divided across the same populations. Though recent studies have involved Asian populations, such as a PheWAS study in the Japanese population ^11^ and construction of an Asian reference genome dataset ^12^, only a few studies have been conducted in Asian populations, and no PheWAS has compared the ethnical differences. Third, in general, the final reports of a PheWAS are mainly comprised of a data-driven analysis and its results, including a multitude of phenotypes and statistical numbers; as a consequence, practical translation of the results to healthcare practices or clinical research has been constricted. While PheWAS incorporates a variety of phenotypes and the associations are provided in a collectively integrated manner that provides good perspective on the holistic view of a system, it is difficult to understand the meaning for particular diseases or phenotypes.

In this study, we addressed these challenges by performing a PheWAS in a Korean population based on the deep phenotyping of a health check-up database. All phenotypes were corroboratively defined after performing laboratory tests, imaging tests, endoscopic tests, and detailed questionnaires on the same day using the same protocols and machines for individual participants. This comprehensive health check-up database merged with a biobank and specific to a Korean population is an unprecedented and unique database, which makes our PheWAS different from those previous. We attempted to compare our PheWAS results with results from the UK Biobank (UKBB) and Biobank Japan project (BBJ). We also leveraged cross-phenotype associations to perform systematic analyses of the PheWAS results, which consist of polygenicity, pleiotropy, a bipartite gene network, and a bipartite phenotype network. To ensure robustness of the PheWAS results, we further dissected them to suggest interpretations applicable to healthcare practice and clinical research (Figure 1).

**Figure 1.**
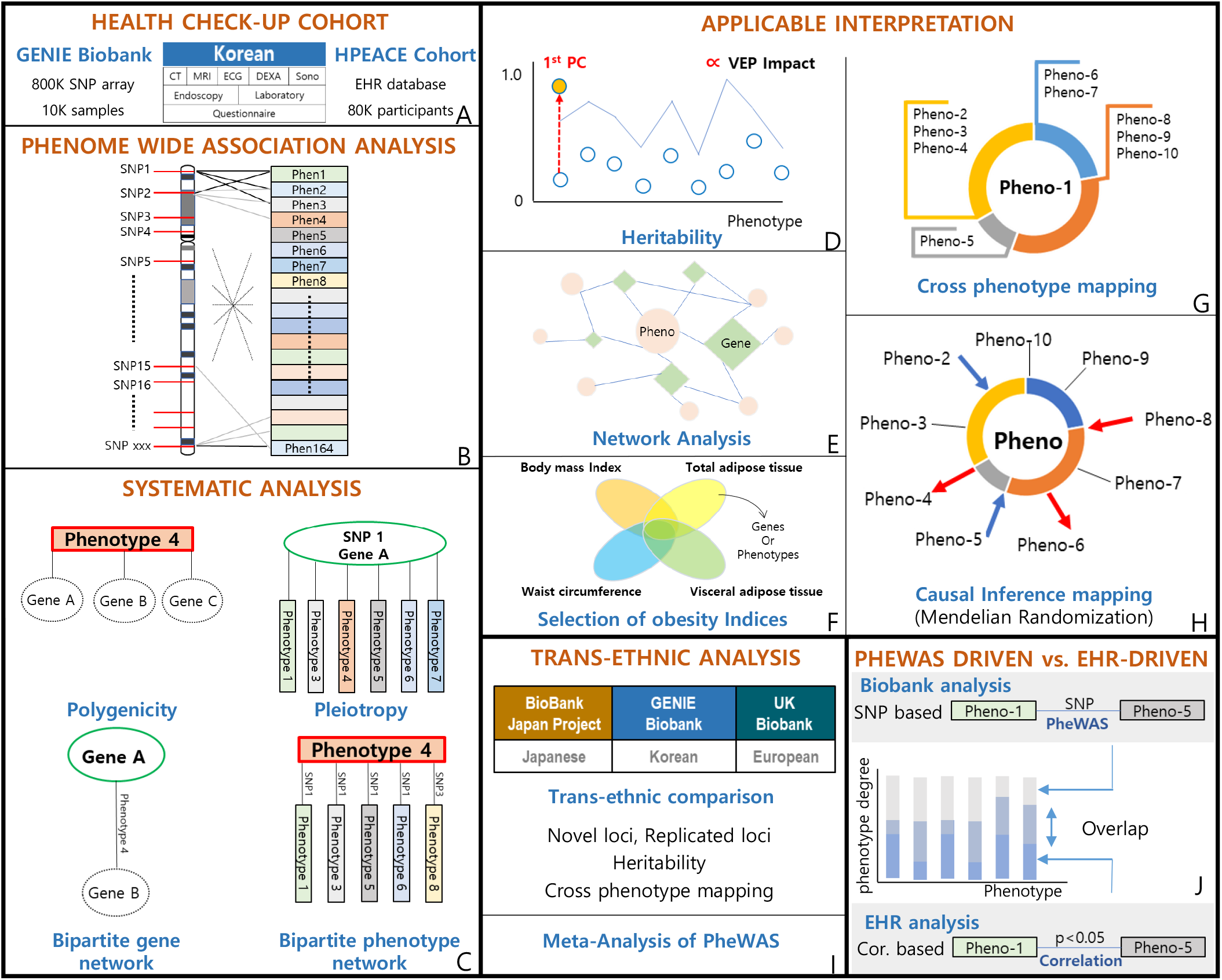
Overview of the study design. A. We utilized a health check-up cohort whose participants had collected CT, MRI, electrocardiography (ECG), bone densitometry (DEXA), ultrasonography (sono), gastroscopy (endoscopy), blood and urine laboratory tests, and questionnaires for medical and medication histories. Sub-cohorts of this cohort are the Gene-EnvironmeNtal IntEraction and phenotype (GENIE) cohort, which includes biobank data from 800K SNP array for 10K samples, and the Health and Prevention EnhAnCEment (H-PEACE) cohort, which includes an EHR database of the health check-up results for 80K participants. B. Phenome wide association study (PheWAS) was performed for 136 phenotypes adjusting for age, sex, and PC1-PC3. C. We leveraged cross-phenotype associations to perform systematic analysis of the PheWAS results, which were polygenicity, pleiotropy, a bipartite gene network, and a bipartite phenotype network. The details are described in Figure 4. D. To ensure robustness of the PheWAS results, we further dissected the results to suggest applicable interpretations. The heritability for each phenotype was calculated using LD regression. We evaluated the correlation between phenotype heritability and the effect of the loci on genes and protein sequences associated with phenotypes, provided by Ensemble variant effect predictor (VEP). We performed a principal component analysis (PCA) for each phenotype using the connections in the bipartite phenotype network, calculated heritability for the 1st principal component (1st PC) of that phenotype, and compared heritability values of the single phenotype vs. its 1st PC. E. Using cross-phenotype association information, we constructed phenotype-phenotype and phenotype-genotype networks to find the hidden relationships among phenotypes or genotypes, as well as to find hub genes or hub phenotypes. F. We visualized the comparison of obesity indices (body mass index, waist circumference, visceral adipose tissue, and total adipose tissue amount) by drawing a Venn diagram of the cross-phenotype associations of phenotypes or genes. G. We constructed cross-phenotype mappings, which have a core phenotype (Pheno-1 in the figure) and branches of connected phenotypes that share loci. These were partitioned by color according to the biological system involved. H. We estimated causal inferences in the phenotype pairs from cross-phenotype associations using Mendelian randomization and constructed a causal inference map. I. Utilizing summary statistics from the UK Biobank and BioBank Japan Project, we performed trans-ethnic and trans-nationality analysis among Korean, European, and Japanese populations. We performed a PheWAS meta-analysis and trans-ethnic comparisons for significant loci, heritability, and cross-phenotype mapping. J. We compared phenotype-phenotype pairs generated from SNP-based cross phenotype-association in the Biobank analysis with those generated from correlation analysis in the EHR-based H-PEACE cohort. We evaluated the overlap or exclusiveness of pairs for each phenotype by phenotype degree.

The results of this work will provide researchers and clinicians with a panoramic overview of the connections among phenotypes, diseases, genes, and loci, allow them to understand healthcare in the perspective of precision medicine, and also let them potentially uncover hidden connections among aspects of health and disease.

## Subjects and Methods

### Gene-Environment of Interaction and phenotype (GENIE) cohort

In this study, we used data from the Gene-Environment of Interaction and phenotype (GENIE) cohort, and the Health and Prevention Enhancement (H-PEACE) cohort, at the Seoul National University Hospital (SNUH) Healthcare System Gangnam Center. GENIE provides a comprehensive database of biomarkers related to non-communicable diseases, lifestyle, medical history, environmental factors, and individual genetic information. The details of the cohort have been described previously ^13^. Briefly, the SNUH Gangnam Center provides comprehensive health check-ups and screening, with nearly 20,000 people visiting this center annually. All participants go through complete questionnaires, physical examinations, laboratory blood and urine tests, abdominal sonography, and gastroscopy. Selectively and on participant request, they also receive advanced tests such as coronary computed tomography (CT), gastroscopy, abdominal CT, and brain magnetic resonance Imaging/magnetic resonance angiography (MRI/MRA). The study population is predominantly Korean. As per consent, we collected blood samples and aliquoted several blood specimens. We also annotated the H-PEACE cohort as an electrical health record (EHR) database of comprehensive health checkups from the Korean population and the GENIE cohort as a genotype database linked to the EHR database. Further preprocessing was conducted for the information in the EHR database. Free text records and questionnaire answers were manually curated by clinicians based on the definitions shown in Table S1. Logical errors and artifacts in the results were manually probed and corrected.

### Ethics statement

The Institutional Review Board (IRB) of the Seoul National University Hospital approved the biorepository with informed consent (IRB number 1103-127357). Construction of the GENIE cohort was approved by the IRB (IRB number H-1505-047-671). We retrospectively collected the clinical and genetic data, for which the IRB approved this study protocol (IRB number 1706-055-858) and waived additional informed consent. The study was performed in accordance with the Declaration of Helsinki.

### Genotype data quality control and imputation

At the time of this study, a total of 10,349 individuals had been genotyped using the Affymetrix Axiom KORV 1.0-96 Array (Thermo Fisher Scientific, Santa Clara, CA, USA) by DNA Link, Inc. This array, referred to as the Korean Chip, was designed by the Center for Genome Science, Korea National Institute of Health; optimized for the Korean population; and available through the K-CHIP consortium. A Korean Chip comprises >833,000 markers, including >247,000 rare-frequency or functional variants estimated from the sequencing data of >2,500 Koreans ^14^.

We performed systematic quality control (QC) on the raw genotype data. SNPs with minor allele frequencies <1%, low marker call rate (<5%), and significant deviation from Hardy-Weinberg equilibrium in controls (HWE < 1e-05) were excluded. Samples with discordant sex info (0.3 < X-chr homozygosity < 0.8, = PROBLEM), low sample call rate (call rate < 0.9, mind 0.1), or extreme heterozygosity (heterozygosity rate > mean +/- 3 SD), along with one individual from any related pairs identified (IBD >= 0.125), were excluded. After quality control was performed, 548,755 SNPs remained. GWAS imputation was carried out using Eagle 2.4.1 (https://data.broadinstitute.org/alkesgroup/Eagle/) and Minimac3 (https://genome.sph.umich.edu/wiki/Minimac3). We used the Northeast Asian Reference Database (NARD) + 1000 Genome Phase 3 database (1KG) re-phased panel as the reference panel and the NARD imputation server (https://nard.macrogen.com) for imputation. NARD ^15^ includes the whole-genome sequencing data of 1,779 individuals from Korea, Mongolia, Japan, China, and Hong Kong, which are not present in 1KG. We compared the imputation quality of chromosome 22 using different reference panels, such as NARD vs. 1KG vs. NARD + 1KG, and determined that the NARD + 1KG panel had the best accuracy (Table S2). Analysis included only high-quality imputed common SNPs, which were those having minor allele frequency >0.01 and imputation R^2^ (Minimac3’s r-squared metric) >0.7. After sample-level QC, genotype-level QC, and imputation, a total of 6,860,342 SNPs from 9,742 individuals were included in the analysis.

The influence of ethnicity was assessed through analysis of population stratification using principal component analysis (PCA) implemented in EIGENSOFT package v6.1.4. We used the first three principal components (PCs) to adjust for population stratification (Figure S2). The steps by which the raw data was preprocessed are shown in Figure S3.

### Phenotype Data

From the comprehensive health check-up database, we manually collated 65 phenotypes as categorical case/control outcomes and 71 phenotypes as continuous numeric outcomes. Tests corroborative of the 136 phenotypes were abdominal/coronary CT scan, brain MRI/MRA, abdominal ultrasonography, esophagogastroduodenoscopy, fundoscopy, tonometry, electrocardiography, bone mineral densitometry (dual-energy x-ray absorptiometry, DEXA), blood/urine test, spinal X-ray, body composition analyzer (InBody^ⓡ^), and questionnaire interview (participant reported phenotypic data). Detailed methods of the test protocols have been described previously ^13^. The phenotypes were systematized into 13 biological categories according to the body system involved: anthropometric measure (AM), cerebro-cardio-vascular (CV), digestive system (DS), endocrine and metabolism (EM), hematologic system (HS), lifestyle (LS), mental and emotional (ME), minerals (MN), musculoskeletal (MC), ophthalmic system (OS), pulmonary system (PS), renal system (RS), and tumor marker (TM). Detailed information on the phenotypes, such as their definitions, categories, associated data formats, and associated tests, are provided as a glossary in Table S1. An overview of the phenotypes is given in Table 1.

**Table 1.** Overview of the studied phenotypes.

| Category | Phenotype | Symbol | Significant loci count<br>($p < 1 \times 10^{-4}$) | Significant loci count<br>($p < 4.916 \times 10^{-10}$) | Significant gene count<br>($p < 1 \times 10^{-4}$) | Heritability<br>($h^2$) |
| --- | --- | --- | --- | --- | --- | --- |
| AM | Anthropometric measure | AM |  |  |  |  |
| AM | Height | Ht | 2415 | 5 | 257 | 0.3221 |
| AM | Weight | Wt | 995 | 0 | 132 | 0.2292 |
| AM | Body mass index | BMI | 886 | 0 | 142 | 0.2375 |
| AM | Skeletal muscle mass | Skeletal_m | 1479 | 0 | 192 | 0.2769 |
| AM | Body fat mass | Fat_mass | 1135 | 0 | 144 | 0.1995 |
| AM | Body fat percent | Fat_percent | 1324 | 0 | 149 | 0.2142 |
| AM | Waist circumference | WC | 841 | 0 | 129 | 0.1781 |
| AM | Total adipose tissue area | TAT | 1251 | 0 | 128 | 0.1505 |
| AM | Visceral adipose tissue area | VAT | 981 | 0 | 131 | 0.1082 |
| CV | Cerebro-cardio-vascular | CV |  |  |  |  |
| CV | Heart rate | HR | 1318 | 40 | 130 | 0.1681 |
| CV | Axis on EKG | Heart_axis | 959 | 0 | 142 | 0.1496 |
| CV | EKG : Sinus bradycardia | Sinusbradycardia | 862 | 40 | 99 | 0.1062 |
| CV | EKG : Right bundle branch block | RBBB | 786 | 0 | 141 | 0 |
| CV | EKG : 1st degree atrioventricular block | 1DAVB | 894 | 0 | 156 | 0.0119 |
| CV | EKG : Myocardial infarction | infarct | 853 | 0 | 160 | 0.0915 |
| CV | EKG : Myocardial ischemia | EKG_ischemia | 1459 | 0 | 276 | 0.2081 |
| CV | Coronary CT : Coronary calcium score | Cal_score | 2688 | 19 | 629 | 0.1278 |
| CV | Coronary CT : Coronary vascular plaque | cor_plaque | 1241 | 0 | 114 | 0 |
| CV | Coronary CT : Coronary vascular stenosis | cor_stenosis | 654 | 0 | 102 | 0 |
| CV | Coronary CT : Aortic dilatation | ao_dilatation | 619 | 0 | 127 | 0.1247 |
| CV | Brain unidentified bright object (UBO) | br_UBO | 519 | 0 | 92 | 0.1272 |
| CV | Brain small vessel disease | br_SVD | 789 | 0 | 117 | 0.0202 |
| CV | Brain vascular atherosclerosis | br_atherosclerosis | 521 | 0 | 105 | 0.1204 |
| CV | Brain vascular stenosis | br_stenosis | 901 | 0 | 182 | 0.1987 |
| CV | Brain aneurysm | br_aneurysm | 720 | 0 | 111 | 0.147 |
| CV | Brain atrophy | br_atrophy | 1246 | 0 | 166 | 0.2294 |
| CV | Diagnosed of hypertension | HTN | 1039 | 0 | 138 | 0.1024 |
| DS | Digestive system | DS |  |  |  |  |
| DS | Gall bladder adenomyomatosis | GB_adenom | 817 | 0 | 140 | 0.0733 |
| DS | Pancreas IPMN | IPMN | 873 | 0 | 164 | 0.0875 |
| DS | Liver hemangioma | Hemangioma | 714 | 4 | 121 | 0.0003 |
| DS | Gall bladder cholecystitis | Cholecyst | 836 | 1 | 156 | 0.0232 |
| DS | Gall bladder stone | GB_stone | 765 | 0 | 135 | 0.0276 |
| DS | Gall bladder polyp | GB_polyp | 904 | 1 | 122 | 0.1163 |
| DS | Fatty liver | Fatty_liver | 849 | 144 | 111 | 0.1332 |
| DS | Atrophic gastritis | AG | 610 | 0 | 103 | 0.015 |
| DS | Intestinal metaplasia of stomach | IM | 1074 | 0 | 151 | 0.1527 |
| DS | Duodenal ulcer | Duodenal ulcer | 833 | 54 | 106 | 0 |
| DS | Gastric ulcer | GU | 1000 | 0 | 200 | 0.0315 |
| DS | Gastroesophageal reflux disease | GERD | 565 | 0 | 101 | 0.0143 |
| DS | Serum total protein | Prot | 945 | 52 | 203 | 0.1993 |
| DS | Serum albumin | Alb | 1310 | 21 | 231 | 0.2325 |
| DS | Serum total bilirubin | Tbil | 2570 | 1151 | 137 | 0.274 |
| DS | Alkaline phosphatase | ALP | 2631 | 299 | 203 | 0.1203 |
| DS | Glutamic oxaloacetic transaminase | GOT | 2209 | 8 | 462 | 0.0334 |
| DS | Glutamic pyruvic transaminase | GPT | 1266 | 6 | 255 | 0.0609 |
| DS | Gamma-Glutamyl Transferase | GGT | 2716 | 78 | 512 | 0.0818 |
| DS | Gastric cancer | Gas_Ca | 982 | 0 | 207 | 0.1719 |
| DS | Hepatitis B virus surface antigen | HBV | 3762 | 324 | 252 | 0.1679 |
| DS | Hepatitis C virus antibody | HCV | 1119 | 0 | 231 | 0.0809 |
| EM | Endocrine and metabolism | EM |  |  |  |  |
| EM | Fasting blood glucose level | glucose | 1842 | 0 | 212 | 0.1116 |
| EM | Uric acid | Urate | 3977 | 1261 | 284 | 0.2186 |
| EM | Triglycerides | TG | 2676 | 333 | 258 | 0.1385 |
| EM | HDL cholesterol | HDL | 2036 | 442 | 171 | 0.2471 |
| EM | Hemoglobin A1c | HBA1C | 1861 | 0 | 245 | 0.1084 |
| EM | Free T4 | T4 | 1652 | 2 | 194 | 0.1547 |
| EM | Thyroid-Stimulating Hormone | TSH | 15064 | 741 | 2549 | 0.1016 |
| EM | Total cholesterol | T_chol | 1061 | 17 | 151 | 0.0678 |
| EM | LDL cholesterol | LDL | 1279 | 63 | 142 | 0.0367 |
| EM | Metabolic syndrome | Metabolic syndrome | 811 | 2 | 132 | 0.1583 |
| EM | Thyroid cancer | Thy_Ca | 836 | 0 | 225 | 0.0023 |
| EM | Breast cancer | Br_Ca | 952 | 0 | 210 | 0.0522 |
| EM | Diagnosed of diabetes | DM | 1507 | 0 | 187 | 0.0824 |
| EM | Diagnosed of dyslipidemia | Dyslipid | 1103 | 2 | 150 | 0.1251 |
| HS | Hematologic system | HS |  |  |  |  |
| HS | White blood cell count | WBC | 1629 | 143 | 153 | 0.1454 |
| HS | Platelet count | PLT | 3040 | 185 | 278 | 0.2375 |
| HS | Neutrophil percent among WBC | Neut | 2080 | 250 | 200 | 0.1423 |
| HS | Lymphocyte percent among WBC | LYMPHO | 1978 | 247 | 190 | 0.1524 |
| HS | Monocyte percent among WBC | MONOCYTE | 1948 | 18 | 194 | 0.2067 |
| HS | Eosinophils percent among WBC | EOSINOPHIL | 3109 | 11 | 343 | 0.2822 |
| HS | Basophils percent among WBC | BASOPHIL | 4043 | 293 | 373 | 0.2941 |
| HS | Red blood cell count | RBC | 1997 | 209 | 181 | 0.2582 |
| HS | Hemoglobin | HB | 1707 | 12 | 199 | 0.1854 |
| HS | Mean corpuscular volume | MCV | 3270 | 250 | 251 | 0.2444 |
| HS | Mean corpuscular hemoglobin | MCH | 3077 | 134 | 358 | 0.2204 |
| HS | Mean corpuscular hemoglobin concentration | MCHC | 4982 | 1266 | 979 | 0.1389 |
| HS | Plateletcrit | PCT | 2747 | 68 | 299 | 0.2023 |
| HS | Mean Platelet Volume | MPV | 3843 | 188 | 591 | 0.1353 |
| HS | Prothrombin time | PT | 4515 | 227 | 846 | 0.061 |
| HS | Activated Partial Thromboplastin Time | aPTT | 2092 | 691 | 181 | 0.1725 |
| HS | Hematocrit | HCT | 962 | 0 | 170 | 0.1544 |
| HS | Red blood cell distribution width | RDW | 2489 | 146 | 244 | 0.1575 |
| LS | Life style | LS |  |  |  |  |
| LS | Smoking history | Smoke | 939 | 0 | 99 | 0.062 |
| LS | Alcohol consumption | Alcohol | 2158 | 612 | 156 | 0.0908 |
| LS | Exercise amount | Exercise | 1657 | 1 | 357 | 0 |
| LS | Education level | education | 558 | 0 | 134 | 0.0264 |
| LS | Marital status | marital | 0 | 0 | 0 | 0.0048 |
| LS | Coffee consumption | Coffee | 680 | 17 | 109 | 0.0317 |
| LS | Nocturia per night | Nocturia | 652 | 0 | 98 | 0.0339 |
| ME | Mental and emotion | ME |  |  |  |  |
| ME | Sleep onset latency | SOL | 503 | 0 | 112 | 0.0931 |
| ME | Wake Time After Sleep Onset | WASO | 869 | 0 | 121 | 0.066 |
| ME | Depressed mood | Depressed | 1043 | 0 | 163 | 0.1041 |
| ME | Appetite change increase | Appetite_inc | 808 | 0 | 114 | 0 |
| ME | Diminished cognitive functioning | Cognitive function | 918 | 0 | 152 | 0.0769 |
| ME | Worthlessness or guilty feeling | Guilty feeling | 1033 | 0 | 104 | 0.0399 |
| ME | Suicidal ideation | Suicidal | 1334 | 0 | 285 | 0.2812 |
| ME | Loss of interest or pleasure | Loss_of_interest | 993 | 0 | 160 | 0.169 |
| ME | Fatigue | Fatigue | 888 | 0 | 158 | 0.046 |
| ME | Psychomotor retardation | retardation | 869 | 0 | 149 | 0.0698 |
| ME | Psychomotor agitation | agitation | 1093 | 0 | 188 | 0.0663 |
| ME | Depression score | depression_score | 839 | 0 | 141 | 0.0614 |
| MN | Minerals | MN |  |  |  |  |
| MN | Calcium level | CALCIUM | 3792 | 597 | 763 | 0.2503 |
| MN | Phosphorus level | PHOSPHORUS | 1461 | 1 | 177 | 0.106 |
| MN | Sodium level | Na | 3992 | 776 | 821 | 0.1384 |
| MN | Potassium level | K | 632 | 0 | 94 | 0 |
| MN | Chloride level | CL | 763 | 3 | 213 | 0.0967 |
| MN | CO2 level | TCO2 | 759 | 0 | 116 | 0.0499 |
| MN | Vitamin D3 | VitD3 | 1271 | 8 | 117 | 0.0743 |
| MC | Musculoskeletal | MC |  |  |  |  |
| MC | Bone density by DEXA | DEXA | 799 | 0 | 88 | 0.2982 |
| MC | Spondylosis | spondylosis | 419 | 0 | 73 | 0 |
| MC | Spondylolisthesis | spondylolisthesis | 939 | 0 | 147 | 0.4245 |
| MC | Compression fracture | compression_fracture | 1189 | 0 | 229 | 0.4589 |
| MC | Intervertebral disc space narrowing | spine_disc_narrowing | 529 | 0 | 97 | 0.0603 |
| OS | Ophthalmic system | OS |  |  |  |  |
| OS | Cataract | cataract | 865 | 0 | 98 | 0.0214 |
| OS | Drusen | drusen | 842 | 0 | 124 | 0 |
| OS | Macular change | macular_change | 881 | 0 | 137 | 0.0347 |
| OS | Optic disc cupping | optic_disc_cupping | 755 | 0 | 133 | 0.0856 |
| OS | Optic nerve fiber loss | optic_fiber_loss | 886 | 0 | 144 | 0.1659 |
| OS | Intraocular pressure, right | IOP_rt | 1468 | 30 | 263 | 0.156 |
| OS | Intraocular pressure, Left | IOP_Lt | 1451 | 4 | 175 | 0.1074 |
| PS | Pulmonary system | PS |  |  |  |  |
| PS | Forced vital capacity (L) | FVC_L | 1519 | 0 | 188 | 0.2408 |
| PS | Forced vital capacity (%) | FVC_percent | 1524 | 0 | 192 | 0.2426 |
| PS | First second of forced expiration (L) | FEV1_L | 1474 | 0 | 182 | 0.2895 |
| PS | First second of forced expiration (%) | FEV1_percent | 2066 | 0 | 168 | 0.2876 |
| PS | FEV1/FVC | FEV1_FVC_percent | 1611 | 86 | 228 | 0.2055 |
| PS | Pulmonary punction test category | PFT_categoric | 563 | 0 | 95 | 0.081 |
| RS | Renal system | RS |  |  |  |  |
| RS | Blood Urea Nitrogen | BUN | 2551 | 123 | 450 | 0.1825 |
| RS | Renal stone | Renal_stone | 824 | 0 | 143 | 0.1145 |
| RS | Creatinine | CREATININE | 2059 | 29 | 399 | 0.2535 |
| RS | Estimated glomerular filtration rate | GFR | 1353 | 34 | 207 | 0.2791 |
| RS | Urine pH | urine_PH | 3432 | 651 | 772 | 0.1166 |
| RS | Urine albumin | urine_PU_cat | 1388 | 0 | 212 | 0.1625 |

| TM | Tumor marker | TM |  |  |  |  |
| --- | --- | --- | --- | --- | --- | --- |
| TM | Cancer Antigen 125 | CA125 | 7923 | 145 | 1508 | 0 |
| TM | Carbohydrate antigen 19-9 | CA19_9 | 27140 | 936 | 4624 | 0.0312 |
| TM | Alpha Fetoprotein | AFP | 4244 | 119 | 654 | 0.1803 |
| TM | Carcinoembryonic antigen | CEA | 1835 | 202 | 375 | 0.0356 |
| TM | Prostate-Specific Antigen | PSA | 12999 | 279 | 2559 | 0.1082 |

### Statistical and computational analyses

#### Phenome-wide association study

We used PLATO ^16^ to run logistic regression analysis on 65 categorical outcomes and linear regression analysis on 71 continuous outcomes, incorporating 6,860,342 genetic variants in an additive model. We included age, sex, and the first three principal components to adjust for any potential confounding bias due to these variables. To identify significant results, we implemented multiple test correction through LD-aware Bonferroni correction. The conventional Bonferroni test assumes that the association tests for all SNPs are independent and thus divides the alpha by the total number of tests. For our study, instead of correcting *p*-values with the total number of SNPs, we use LD pruning to identify independent SNPs ^17^. The threshold we used for association between SNPs was r^2^ = 0.3, which is provided by Sobota et al. for the East Asian population ^18^. We established genome-wide significance at *P* < 4.92×10^−10^.

Further exploratory analyses were performed using the associated 260,923 loci with a less stringent *P* < 1 x 10^−4^. Though we used the LD pruning method for Bonferroni correction, the *p*-value was still stringent. Thus, in addition to analyzing associations with a stringent *p*-value cutoff, this exploratory threshold allowed us to further expand the boundaries of research by involving a much wider PheWAS landscape ^17^.

To perform systematic analysis of the PheWAS results, we leveraged cross-phenotype associations, in which one locus is associated with multiple phenotypes ^19^. Such associations include polygenic inheritance, where a phenotype is influenced by more than one gene ^20^ (Figure S4A); and pleiotropy, where a locus or a gene affects more than one phenotype ^21^ (Figure S4B). To further explore and understand polygenicity and pleiotropy, we constructed two networks: a bipartite phenotype network, connecting phenotypes that shared at least one locus ^19^ (Figure S4C) and a bipartite gene network, connecting genes that shared at least one phenotype ^19^ (Figure S4D). In these connections or networks, the degree property indicates the number of direct connections between one core component and other components. For each core gene/phenotype, the number of genes associated or connected with it is defined as its gene degree, and the number of phenotypes associated or connected is its phenotype degree.

We used the cross-phenotype association information to construct a phenotype-phenotype network and a phenotype-genotype network in order to find hidden relationships among phenotypes or genotypes and also to identify hub genes or hub phenotypes. The Gephi software (https://gephi.org/) was used to visualize the network ^22^.

#### Functional Annotations (p value < 1 x 10^−4^)

We mapped genetic associations using the Ensembl Variant Effect Predictor (VEP) ^23^ in order to annotate the functional relevance of significant loci. Using the VEP annotation, we classified the biological consequences of loci in coding regions (stop-gained variant, slice acceptor variant, splice donor variant, and missense variant) and in non-coding regions. We also annotated UKBB and BBJ variants with VEP to conduct trans-ethnic and trans-national comparisons as described in a later section.

#### Estimated heritability

To determine the contributions of genetic variants to the risk of certain phenotypes, we estimated the heritability of each phenotype. We estimated heritability using LD Score regression with LDSC (version 1.0.1) ^24^ on summary statistics from the PheWAS for all phenotypes. For this analysis, we used the East Asian LD Scores from 1000 Genomes as reference LD Score, which served as the independent variable in the LD Score regression (ref-ld-chr) and regression weights (w-ld-chr). General instructions and the East Asian LD Scores from 1000 Genomes are provided here: https://github.com/bulik/ldsc.

Since there are issues with missing heritability when estimating from genotype data ^25^, we utilized the reduction of phenotype dimensions in the cross-phenotype associations to improve explanation of the amount of heritability. We generated a heritability estimation model using the bipartite phenotype network explored by the pleiotropic nature of variants; this model incorporates all phenotypes in connection with a given phenotype through principal component analysis (PCA). Then, we compared the sole heritability of the given phenotype vs. the PCA-based heritability of the same phenotype. To do this, we first separately imputed continuous and discrete phenotypes using the *mice* package in R ^26^. Then, we used PCA to generate principal components (PCs) and reduce the information of phenotypes sharing variants with the phenotype of interest. Next, we estimated the heritability of each phenotype using the generated PCs. If the group consisted solely of continuous phenotypes, then we simply used the *prcomp* package in R to calculate the PCs. However, if the phenotypes in the group were mixed, i.e. discrete and continuous, then we used the *PCAmixdata* package in R. We took the first principal component for each group, conducted a genome-wide association analysis, and estimated heritability using the summary statistics generated by LDSC as described above.

#### Comparison in different populations

To compare results across diverse populations, we performed a trans-ethnic comparison utilizing PheWAS results from a European population and a trans-national comparison utilizing results from a Japanese population. For European population, data from the UK Biobank (UKBB) ^27^ was used; for the Japanese population, data from the Biobank Japan Project (BBJ) ^11^ was used. We downloaded the summary statistics and estimated heritability results of the phenotypes of these results from the following URLs: http://www.nealelab.is/uk-biobank/ and http://jenger.riken.jp/en/result. We tabulated lists of the phenotypes in the UKBB and BBJ and searched for those that were most similar to phenotypes in our database. The manually curated overlapping phenotypes among GENIE, UKBB, and BBJ are given in Table S3.

#### Mendelian randomization

To better understand the causal inferences in cross phenotype mapping, we performed Mendelian randomization (MR) analysis on the phenotype pairs connected in the bipartite phenotype network. To avoid potential bias due to sample overlap between exposure and outcome, we split our dataset into two equal sets by random assignment of samples. PheWAS was conducted on each dataset separately to generate the summary statistics that were used as input to MR. Additionally, significant SNPs (*P* < 1×10^−4^) from the initial PheWAS with all samples were used as instrument variables (IV). Furthermore, all IVs that were significant in outcome (*P* < 0.01) were removed, as IVs should not be directly associated with outcome. We calculated *p*-values using the inverse-variance weighted (IVW) method from the *MendelianRandomization* package in R ^28^. We adjusted for multiple testing using FDR correction. We also performed sensitivity analysis using MR-egger and the median-based method.

#### Meta-analysis of PheWAS

We performed meta-analysis using our PheWAS results and the BBJ results for all phenotypes that were available in both datasets. The BBJ summary statistics came from different studies, requiring harmonization of the files. Phenotype matches between GENIE and BBJ are listed in Table S3. Some of the phenotypes from GENIE matched to multiple phenotypes in BBJ; in such cases, we carried out meta-analysis separately for each BBJ phenotype. The meta-analysis was implemented using METAL ^29^. The overall scheme of our study is shown in Figure 1.

## Results

After QC, the study population of the GENIE cohort included 9,742 participants, comprising 5,696 males and 4,046 females, with average age 50.7 +/- 10.0 years. The characteristics of the study population are given in Table S4.

### Phenome-wide association analysis for 136 phenotypes

#### GENIE cohort (Korean population)

From the PheWAS on 136 phenotypes, we found significant associations for 65 phenotypes (50 from continuous variables, 13 from categorical variables) and 14,101 SNPs at *P* <= 4.92×10^−10^. The counts of significant loci and genes associated with each phenotype are given in Table S5. Among continuous phenotypes, the top five most significant were activated partial thromboplastin time (aPTT), LDL cholesterol, serum total bilirubin, uric acid, and carcinoembryonic antigen (CEA). Among categorical phenotypes, the top five most significant were alcohol consumption, fatty liver, duodenal ulcer, coffee consumption, and Hepatitis B virus surface antigen (Table S6, Figure S5). In the Manhattan plot, aPTT had two top signal loci, in chromosome 5 (rs17876032, *P* = 3.29×10^−83^) and chromosome 9 (rs676996, *P* = 8.99×10^−72^). Table S7 lists the top five signals from the GWAS results for each phenotype.

We further performed functional annotation for 221,462 unique loci passing the less stringent *p*-value threshold of 1×10^−4^ using Ensembl Variant Effect Predictor (VEP) ^23^. Approximately 1% of variants were in coding regions and 98.885% were in non-coding regions (Figure S6), which result is similar to other large-scale PheWAS ^17^. In coding regions, this annotation identified 22 stop-gained variants, six splice acceptor variants, ten splice donor variants, and 1103 missense variants (Table S8 and S9).

Among the 22 stop-gained variants, we replicated an association between rs121907892 and uric acid that is a well-reported finding unique to the east Asian population (EAS) ^30^, including Koreans ^31^. We further identified the stop-gained variant rs200340875 to be significantly associated with blood urea nitrogen (BUN; *P* = 2.75×10^−07^, beta = −0.373), calcium (*P* = 7.15×10^−08^, beta = −0.044), glutamic oxaloacetic transaminase (GOT; *P* = 9.65×10^−6^, beta = 1.189), mean corpuscular hemoglobin (MCH; *P* = 4.93×10^−7^, beta = 0.21), mean corpuscular hemoglobin concentration (MCHC; *P* = 8.53×10^−16^, beta = 0.228), mean platelet volume (MPV; *P* = 8.06×10^−7^, beta = −0.088), urine protein (*P* = 1.16×10^−10^, beta = −0.135), and sodium (Na; *P* = 1.92×10^−15^, beta = −0.371). According to 1000 Genomes, the minor allele of rs200340875 is reported in African populations (AFR) but not in EAS. The locus of this variant is associated with CD109 Molecule (*CD109*), which has been previously reported with diffuse large B-cell lymphoma ^32^, psoriasis ^33^, and gallbladder malignancy ^34^. Another stop-gained variant identified was rs145035679, which showed a protective effect for CEA (*P* = 1.07×10^−8^, beta = −0.166) and increased risk for carbohydrate antigen 19-9 (CA 19-9; *P* = 5.81×10^−5^, beta = 2.178); this variant is associated with Fucosyltransferase 6 (*FUT6*). *FUT6* has previously reported associations with pancreatic cancer ^35^, breast cancer ^36^, and colorectal cancer ^37^. Among splice acceptor variants, rs112911835 was significantly associated with prothrombin time (PT; *P* = 1.12×10^−10^, beta = − 0.031), while rs112911835 was associated with Long Intergenic Non-Protein Coding RNA 1933 (*LINC01933*), which has no known relationship with disease as of yet. The splice donor variant rs140944893 showed significant association with coronary vessel calcium scoring (*P* = 4.29×10^−5^, beta = −127.7), and is associated with Phospholipase D3 (*PLD3*). This gene is reported to be related to Alzheimer’s disease in EAS ^38^ and EUR ^39^.

#### Comparison with BBJ (Japanese) and UKBB (European)

We systematically compared the significant associations of loci and their genes with phenotypes (*P* < 1×10^−4^) to results from the BBJ and UKBB to determine if our results were replicated in other populations and also to look for novel findings. Originally, each population used a different SNP array platform and a variety of different phenotypes. Accordingly, we first filtered the examined loci and phenotypes to determine overlap with our data, then identified replicated and novel genes and loci. The schematic structures of the trans-ethnical and transnationality comparisons are shown in Figure S7. We identified 52 phenotypes overlapping the Japanese biobank results (42 phenotypes had replicated loci) and 101 phenotypes overlapping with the UK Biobank results (59 phenotypes had replicated loci). Gene-level and locus-level comparisons are respectively given in Table S10 and Table S11.

In the comparison between Korean and Japanese populations, aPTT and serum total bilirubin had high overlap of significant loci. Among the 4,016 loci significantly associated with aPTT in Koreans or Japanese, 920 loci (22.91%) were significant in both; among the 6,159 loci associated with total bilirubin in those populations, 1,263 (20.51%) likewise overlapped. Notably, loci associated with the ophthalmic system (cataract and optic fiber loss), cerebrovascular system (brain stenosis, aneurysm, and atherosclerosis), smoking habit, hepatitis C virus antibody, renal stone, gastric cancer, and bone mineral density were mutually exclusive between Koreans and Japanese.

In the comparison between Korean and UK populations, fewer overlapping loci were identified, with the highest overlap ratio being 9.15% in fatty liver disease; 42 phenotypes did not have any overlap (Figure 2, Figure S8).

**Figure 2.**
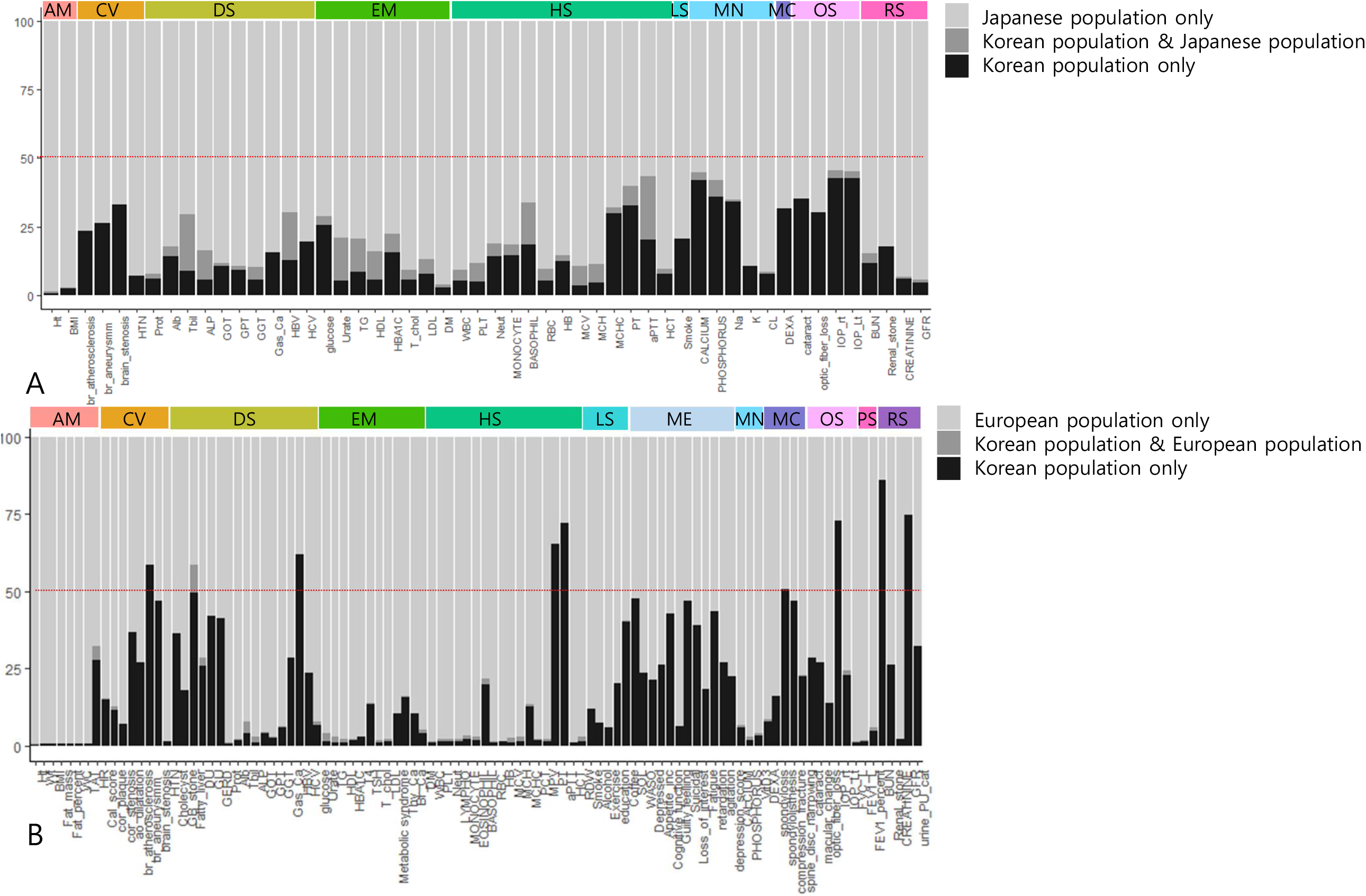
Trans-ethnic, trans-nationality comparison of PheWAS. We compared PheWAS results among Korean, Japanese, and European populations. Phenotypes existing in all datasets were used. We evaluated loci significantly associated only in Koreans (black bar), in both populations (gray bar), and only in the other population (bright gray bar). The colored bar at the top indicates phenotype categories. The Y axis denotes the ratio (%) of loci in each classification, with 100% being the total significant in the compared populations. A. PheWAS result comparison between Korean and Japanese populations. B. PheWAS result comparison between Korean and European populations.

Population comparisons were further investigated for body mass index (BMI) in particular. For this phenotype, 136 loci (0.42% of significant loci) were replicated in the Japanese population and 105 loci (0.07%) in the European population, respectively. Our population showed 583 exclusive loci (1.82%) when compared to the Japanese population, and 669 exclusive loci (0.45%) when compared to the European population. We then looked more deeply into the BMI genes unique to the Korean population. Relative to the Japanese population, 73 genes were exclusively associated with the Korean population; meanwhile, relative to the European population, 53 genes were exclusively associated with the Korean population. Of these genes, 34 (714 loci) were unique relative to both Japanese and European populations (Table S12, Figure S9). Of those unique genes, 23 have previously reported associations with obesity or body weight; the corresponding literature review and references are given in Table S13. The other 11 genes have not been previously reported as associated with obesity in humans, and could be candidate novel genes for BMI or obesity; these were Vesicle Amine Transport 1-Like (*VAT1L*), Uromodulin-like 1 (*UMODL1*), Telomeric Repeat-Binding Factor 2-Interacting Protein 1 (*TERF2IP*), Proline-rich Transmembrane Protein 3 (*PRRT3*), PRRT3 Antisense RNA 1 (*PRRT3-AS1*), Long Intergenic Non-protein Coding RNA 578 (*LINC00578*), Family with Sequence Similarity 225 Member B (*FAM225B*), Cation Channel Sperm-associated 1 (*CATSPER1*), Barrier To Autointegration Factor 1 (*BANF1*), Attractin-Like Protein 1 (*ATRNL1*), and Adherens Junctions-associated Protein 1 (*AJAP1*). Among those genes, *TERF2IP* is known from a mouse study to have roles in regulating adipose function and excess fat accumulation, and also protecting against obesity ^40^. *ATRNL1* has no previous report related to obesity, but Attractin (*ATRN*) has similarity with the mouse mahogany protein, which is involved in controlling obesity ^41,42^. *BANF1* has no known direct association for obesity, but it is reported to suppress expression of S100 calcium-binding protein A9 (*S100A9*) ^43^, which is a candidate marker for obesity in non-type 2 diabetes mellitus ^44^.

### Systematic analysis of the PheWAS results

#### GENIE cohort (Korean population)

To perform a systematic analysis of the PheWAS results, we leveraged cross-phenotype associations, where one locus is significantly associated with multiple phenotypes. For this analysis, significant loci were filtered by a less-stringent threshold, *P* < 1×10^−4^ (loci count = 260,922, gene count = 14,907). The schematic structure for this analysis is shown in Figure S4. Briefly, we constructed: “Possible polygenicity”, in which a phenotype is influenced by more than one gene (Figure S4A, Table S14); “Possible pleiotropy”, in which a locus or gene affects multiple phenotypes (Figure S4B, Table S15); a “bipartite phenotype network” based on the connections among phenotypes sharing at least one locus (Figure S4C, Table S16); and a “bipartite gene network” as the connections among genes shared by at least one phenotype (Figure S4D).

Of the 260,922 significant PheWAS loci, the bipartite phenotype network comprised 23,580 loci (2,902 genes) with 135 phenotypes. There were 1,926 distinct pairs of phenotypes. We calculated the degree properties of core phenotypes in this network (Table S17), where core phenotypes were those nodes connected to several phenotypes by shared variants; an example is phenotype 4 in Figure S4C. Notably, phenotypes in the tumor markers category had relatively high degree of phenotype connection. The highest phenotype degree was obtained for a representative tumor marker for pancreas cancer, *CA 19-9*, with 110 phenotypes connected through sharing of significant loci. Meanwhile, the highest possible polygenicity was observed for mean corpuscular hemoglobin concentration (MCHC), with 782 genes.

The bipartite gene network comprised 14,907 genes, which were connected through sharing associations with the same phenotypes. Table S18 give the gene degree and phenotype degree values for this network. The three genes with the highest phenotype degrees were; CUB and Sushi Multiple Domains 1 Protein (*CSMD1*), RNA-binding Fox-1 Homolog 1 (*RBFOX1*), and Protein Tyrosine Phosphatase Receptor Type D (*PTPRD*); this could be due to possible pleiotropy. The same three genes had the highest gene degree values; gene degree comprises the edges in bipartite gene networks. Notably, *CSMD1* was significantly associated with 58 phenotypes (showing possible pleiotropy) and connected to 12,602 genes through common associations with phenotypes. *CSMD1* has been reported to function as a complement control protein ^45^; complement is implicated in many diseases through the mechanisms of inflammation and autoimmunity ^46^. In some cancers, it functions as a tumor suppressor gene ^47,48^.

#### Bipartite phenotype network comparison with BBJ and UKBB

We compared the bipartite phenotype networks of the GENIE (Korea), BBJ (Japanese), and UKBB (European) cohorts. There were 49 GENIE phenotypes in common among all datasets, which were used to generate the bipartite phenotype network. Figure S10 shows a Venn diagram of the phenotype-phenotype pairs observed in each population; 288 pairs were simultaneously observed in all three populations (Table S19). Notably, these included the pairing of red blood cell count (RBC) and brain vascular atherosclerosis. There are reports of RBC having relation to coronary artery disease ^49^ and stroke mortality ^50^, but not directly to brain vascular atherosclerosis.

##### Applicable interpretation of the PheWAS results

#### Heritability analysis

Heritability was calculated for each of the 136 phenotypes by regression of LD scores (Table S20). The top heritability values were obtained for compression fracture (h2 = 0.459), spondylolisthesis (h2 =0.425), height (h2 = 0.322), and bone mineral density (h2 = 0.298). In terms of biological categories and body systems, the highest heritability values were obtained for the musculoskeletal system (mean h2 = 0.244), the pulmonary system (mean h2 = 0.225), anthropometric measures (mean h2 = 0.213), and the hematologic system (mean h2 = 0.188) (Table S21).

For further interpretation of heritability, we produced an integrated phenotypic trait using a dimensionality reduction approach. Specifically, we performed principal component analysis (PCA) for each phenotype using connected phenotypes from the bipartite phenotype network (for example, phenotype 4 group in Figure S4C). Then, we calculated heritability for the resulting new phenotypic trait, the 1st principal component (1st PC) of the given phenotype. To observe the effect of this dimensionality reduction approach, delta h^2^ was calculated by subtracting the original heritability (mean heritability or heritability of a single phenotype) from the heritability of the 1st PC (Δh^2^ for 1st PC). There were 135 phenotypes that have at least one phenotype in connection by sharing variants. The results are given in Figure S11A and Table S20. Of our 135 phenotypes, 82 had improved heritability values. As shown in Figure S11A, particularly high delta h^2^ values were obtained for spondylosis, exercise, liver hemangioma, and gastroesophageal reflux.

The Ensembl variant effect predictor (VEP) provides information regarding the effect of loci on genes and protein sequences, categorized as modifier impact (usually for non-coding variants or variants affecting non-coding genes, where predictions are difficult or there is no evidence of impact), low impact (mostly harmless or unlikely to change protein behavior), moderate impact (non-disruptive variant that might change protein effectiveness), and high impact (high, disruptive impact on the protein) (https://useast.ensembl.org/Help/Glossary?id=535). We divided the significant loci (1×10^−4^) into two groups according to their annotated impacts, namely “modifier low” vs. “moderate, high”, and evaluated the correlation between impact group and heritability in each phenotype. A significant correlation was observed (*P* = 0.001, correlation (r) = 0.281, 95% CI = 0.117-0.429).

We further compared the heritability in our population with that in the Japanese and European populations (Table S20). Of phenotypes that overlapped with ours, BBJ provides heritability for 35 and UKBB for 101. Since the provided heritability values were determined using different loci and methods, we normalized the heritability to make it comparable. Comparisons to each of the Japanese and UK populations are shown in Figure S12, while the three-way comparison among Korean, Japanese, and UK populations is shown in Figure S11B (33 phenotypes overlapped among the three populations). Generally, most phenotypes had similar trends in heritability across populations. Noticeable differences were observed in percent eosinophils among white blood cell counts (BASOPHIL), prothrombin time in the international normalized ratio (PT), and activated partial thromboplastin time (aPTT). BASOPHIL had relatively high heritability in the Korean population relative to both others. Meanwhile, PT and aPTT, which are biomarkers of coagulation function, showed similar trends in the Korean and Japanese populations, but manifested relatively high heritability in Koreans relative to the UK population.

#### Network analysis

Using cross-phenotype association information, we constructed phenotype-phenotype and phenotype-genotype networks in order to find hidden relationships among phenotypes or genotypes and to discover hub genes or phenotypes.

First, a network representation of gene-phenotype associations related to metabolic syndrome was constructed (Figure 3A). We selected the nodes by filtering for genes associated with metabolic syndrome, which were identified by annotating the significant loci (*P* < 10-4). Then, we filtered for phenotypes significantly associated with those selected genes. In the process, edges corresponding to loci not annotated by VEP were not included. Ultimately, 132 genes associated with metabolic syndrome and 128 phenotypes sharing 102 genes with metabolic syndrome were used to construct the network (Figure 3A). The nodes were colored with respect to gene and phenotype, while the edges are associations between phenotypes and genes. In the metabolic syndrome sub-network, five genes had high degrees of connection and could be considered hub genes: *PTPRD*, DCC Netrin 1 Receptor (*DCC*), Proprotein Convertase Subtilisin/kexin Type 6 (*PCSK6*), Unc-13 Homolog C (*UNC13C*), and Contactin 4 (*CNTN4*). The phenotypes in this network comprised: of cardiovascular diseases, of metabolic diseases, used as markers for obesity, and other various disease. The phenotype nodes included triglyceride (TG), HDL cholesterol (HDL), hypertension, diabetes, and waist circumference (WC). These results give a genetic rationale for the definition of metabolic syndrome in the PheWAS perspective.

**Figure 3.**
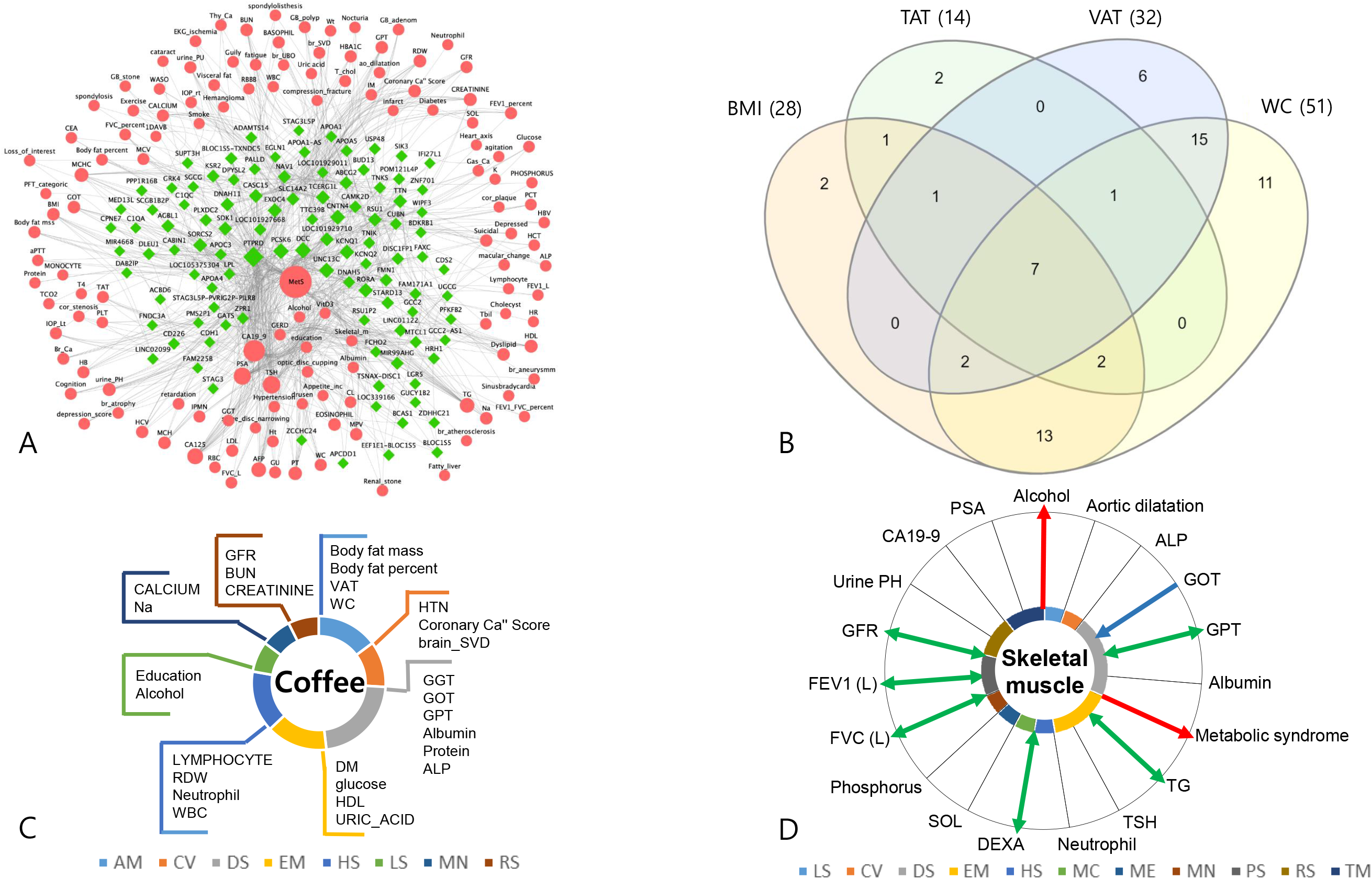
Applicable interpretations. A. **Network analysis** A network representation of gene-phenotype associations related to metabolic syndrome was constructed from 102 genes associated with metabolic syndrome and 128 phenotypes sharing those genes. Each edge is a phenotype-gene association, with genes for significant loci (*P* < 10^−4^) being annotated by VEP. Node size is proportional to degree, which is the number of connections. Pink nodes correspond to phenotypes and green nodes to genes. The genes *PTPRD*, *DCC*, and *PCSK6* had high degree, which implies the possibility of being hub genes. Among phenotype nodes, CA19-9 and TSH were of high degree. B. **Relationships among obesity indices** We visualized the comparison among the obesity indices such as body mass index (BMI), waist circumference (WC), visceral adipose tissue (VAT) and total adipose tissue (TAT) amount by drawing a the venn-diagram for cross phenotype association of phenotypes or genes. There were 7 phenotypes (CA 19-9, body fat percent, body fat mass, GOT, GPT, weight, and metabolic syndrome), which are simultaneously associated with BMI, WC, TAT and VAT. 15 phenotypes (alcohol consumption, coffee consumption, ALP, GGT, creatinine, GFR, glucose, diabetes, HDL cholesterol, hypertension, WBC, neutrophil, RDW, protein and albumin) were associated exclusively with VAT and WC and this intersection had 2 genes associated. C. **Cross-phenotype mapping** Cross-phenotype mappings were generated based on the bipartite phenotype network, in turn constructed from the connections among phenotypes sharing at least one locus. Coffee consumption, which is one of the lifestyle phenotypes, had 31 phenotype degrees in the bipartite phenotype network. D. **Causal inference mapping** We estimated causal inferences in phenotype pairs based on cross-phenotype associations using Mendelian randomization (MR), and constructed a causal inference map. This figure shows the map for skeletal muscle mass (Skeletal muscle). The network grid is based on information from the bipartite phenotype network of Skeletal muscle. We excluded pairs whose biological categories are anthropometric measurements, the same as skeletal muscle. We then performed pair-wise Mendelian randomization for each phenotype pair. Pairs with significant *P* values (false discovery rate [FDR] less than 0.05) are indicated by arrows. The direction of the arrow is the causality result from MR (Blue arrows, WBC as outcome; Red arrows, WBC as exposure; Green arrows, bidirectional). Pairs observed in the bipartite phenotype network but insignificant in MR have straight black lines without arrows.

**Figure 4.**
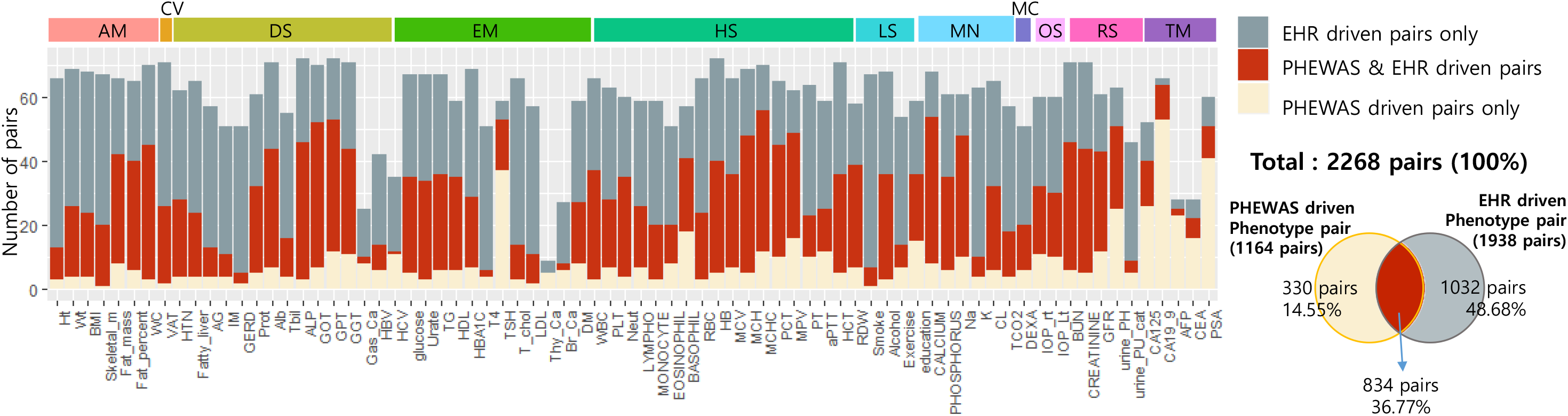
Comparison of phenotype-phenotype pairs between PheWAS driven and EHR-driven analysis. There were 76 phenotypes also recorded in the EHR-driven cohort (H-PEACE cohort). PheWAS-driven pairs (1164) were based on shared SNPs with association *P* < 1×10^−4^, and EHR-driven pairs (1938) on correlation analysis with multi-test corrected *P* < 0.05 (Table S26). Skeletal muscle mass (95%) and alkaline phosphatase (93.48%) had high ratios of overlap, while thyroid cancer (0%) and alpha fetoprotein (8%) had low ratios. In terms of biological categories, the average replication % was highest for anthropometric measurement (86.43%) Of the 1164 pairs from the PheWAS-driven approach, 834 (71.65%) also manifested significance in the EHR-driven analysis.

We also constructed a phenotype-phenotype network using 1,926 phenotype pairs based on shared loci (*P* < 1×10^−4^). Figure S13 shows the phenotype-phenotype network for the whole dataset, and an interactive visualization tool of the phenotype-phenotype network is available (https://hdpm.biomedinfolab.com/genie/).

#### Relationships among obesity indices

Ever since the American Medical Association (AMA) declared obesity to be a disease, interest in and research into obesity has been growing ^51,52^. However, definitions of pathological obesity make inconsistent use of variable traits such as body mass index (BMI), waist circumference (WC), total adipose tissue area (TAT), and visceral adipose tissue area (VAT). There are reports of an obesity paradox when defining obesity by BMI ^53^. The defining parameter for obesity also varies between researchers and with respect to the target disease. To investigate the relationships among these parameters, we constructed Venn diagrams ^54^ and visualized the overlap or exclusiveness among BMI, WC, TAT, and VAT based on the bipartite phenotype network (phenotype level) and pleiotropy/polygenicity potential of genes (gene level). As shown in Figure 3B, connections were observed as quadrant intersections among BMI, WC, TAT, and VAT for seven phenotypes: CA19-9, GOT, GPT, body fat mass, body fat percent, weight, and metabolic syndrome. There were 15 phenotypes connected exclusively with VAT and WC, and the intersection between these traits had two exclusive genes associated. Of the 15 phenotypes, most were crucial intermediate phenotypes that link obesity with diseases. Accordingly, it can be postulated that when defining obesity, VAT or WC would better represent the characteristics of pathogenic obesity. The two genes that are exclusively overlapped between VAT and WC (Figure S14) could be candidate genes for explaining the pathogenic role of obesity. The elements in each set are listed in Table S22.

#### Cross-phenotype mapping

Cross-phenotype mappings were generated based on the bipartite phenotype network, in which the connected phenotypes shared at least one locus.

First, we constructed a cross-phenotype mapping focused on tumor markers. Several tumor markers are used in screening for cancer, monitoring its recurrence, and evaluating its response to interventions. Commonly-used tumor markers include carcinoembryonic antigen (CEA), carbohydrate antigen 19-9 (CA19-9), alpha fetoprotein (AFP), and prostate-specific antigen (PSA); specifically, CEA serves as a marker for colorectal cancer ^55^, CA19-9 for pancreato-biliary cancer ^56^, AFP for liver cancer ^57^, and PSA for prostate cancer ^58^. However, the limitation of using the tumor markers is that it can have low sensitivity or specificity ^59^, such that a test result could be associated with or affected by various non-malignant conditions. For instance, CEA is known to be affected by hemoglobin level ^60^, and CA19-9 is reported to be elevated in nonmalignant respiratory disease ^61^. Table S23 shows the respective connected phenotypes we obtained for tumor markers; among these, CEA is associated with hemoglobin level and CA19-9 with pulmonary function test, which are consistent with previous reports ^60,61^. Figure S15 shows the cross-phenotype mapping for CEA, which could be considered during oncological practice in order to take into consideration all the possible effects of phenotypes other than colorectal cancer progression itself.

Second, we constructed a cross-phenotype mapping focused on lifestyle factors. The analyzed phenotypes included lifestyle factors such as coffee consumption and alcohol consumption. Several studies have shown genotype x environment interactions (G x E) in smoking behaviors ^62^. In this study, we visualized the cross-phenotype mapping for the coffee consumption as a starting point for G x E study in this phenotype. Coffee consumption had 27 phenotypes connected through sharing of significant loci (Figure 3C). Several reports have documented relationships between coffee consumption and obesity ^63^, hypertension ^64^, diabetes ^65^, renal function ^66^, and lipid metabolism ^67^. The phenotypes connected with coffee consumption in this mapping (Table S16) support the previous reports of clinical association studies. In the mapping for alcohol consumption, 38 phenotypes shared significant loci. Various studies have identified heavy alcohol consumption as a risk factor for renal disease ^68^ and coronary artery calcification ^69^. The results of these and other cross-phenotype mappings could provide the genetic background to explain interactions between environmental factors and disease, and might further provide basic knowledge necessary to conduct G x E analysis.

#### Mendelian randomization analysis

We estimated the causal inferences in phenotype pairs based on cross-phenotype associations using Mendelian randomization (MR). Table S24 shows the MR results for each pair having false discovery rate (FDR) < 0.05. Of the phenotype pairs, significant in the cross-phenotype association, 1766 retained significant association after the Mendelian randomization analysis. As shown in Figure 3D, we drew a causal inference mapping centered on skeletal muscle mass. The network grid is based on information from the bipartite phenotype network of skeletal muscle mass. We excluded those pairs whose biological categories were anthropometric measurements, which category includes skeletal muscle mass. The Mendelian randomization analysis yielded nine significant phenotypes, of which one was causal for skeletal muscle mass, two phenotypes were outcomes from skeletal muscle mass, and six had bidirectional relationships with skeletal muscle mass. This analysis revealed that skeletal muscle mass was a significant causal factor for metabolic syndrome and alcohol consumption. Its bidirectional relationships were with bone mineral density, liver function (GPT), pulmonary function (FVC, FEV1), renal function (glomerular filtration rate), and triglyceride.

We also performed Mendelian randomization with a focus on lifestyle factors that were causal exposures in cross-phenotype associations, such as alcohol consumption, coffee consumption, exercise amount, and smoking history. Table S25 shows the significant outcome phenotypes (FDR < 0.05) from this analysis. Alcohol consumption was a significant causal exposure for ten phenotypes, coffee consumption for three phenotypes, exercise amount for six phenotypes, and smoking history for two phenotypes. Coffee consumption was also a significant causal exposure for three anthropometric measurements: body fat mass, visceral adipose tissue area, and waist circumference.

### Comparison of the phenotype-phenotype pairs between PheWAS-driven vs. EHR-driven

“Penetrance” in genetics is the proportion of those individuals carrying a certain genetic variant who also exhibit the associated phenotype, while “expressivity” measures the proportion of individuals that are carriers of a certain variant and show the associated phenotype to a certain extent ^70^. As an indirect method to investigate the penetrance or expressivity of the significant loci identified in our study, we repeated bipartite phenotype network construction using an electronic health records (EHR)-driven method. This clinical database consisted of 81,086 distinct participants who went through comprehensive health check-ups from 2004 to 2015 in the SNUH Gangnam Center (H-PEACE cohort). The tests and questionnaires included most of the phenotypes used in the PheWAS study; specifically, 76 phenotypes were also recorded for this cohort. PheWAS-driven pairs (1164 pairs) were selected based on shared SNPs with association *P* < 1×10^−4^, and EHR-driven pairs (1938 pairs) were selected based on correlation analysis with multi-test corrected *P* < 0.05. We compared these phenotype-phenotype pairs (Table S26) and evaluated the overlap or exclusiveness of the pairs for each phenotype. Of the 1164 pairs identified in the PheWAS-driven approach, 834 (71.65%) also manifested significance in the EHR-driven analysis. As shown in Figure 5 and Table S27, high ratios of overlap were identified for skeletal muscle mass (95%) and alkaline phosphatase (93.48%), and low ratios for thyroid cancer (0%) and alpha fetoprotein (8%). When viewed in terms of biological category, the highest average % replication was obtained for anthropometric measurement (86.43%).

#### Meta-analysis of PheWAS from Korean and Japanese populations

We performed a PheWAS meta-analysis by incorporating our data with the BBJ data (Japanese population). The results are given in Table S28, Figure S16 and Figure S17. All 51 phenotypes used in the meta-analysis had an increased number of significant variants in the Korean population, while 37 phenotypes had variants uniquely significant in the meta-analysis. Furthermore, height, diabetes and body mass index had more than 100 variants that were uniquely identified as significant in the meta-analysis.

## Discussion

With the advancements in healthcare research that are being driven by big data, increasing efforts are being made to carry out data-wide association studies. PheWAS is one of the tools in that paradigm. However, previous studies faced major challenges in terms of deep phenotyping due to generally using ICD codes, which have limited clarity in their definitions; making the results applicable and interpretable in healthcare practices and clinical research; and the characteristics of population genetics, being highly affected by race and ethnicity. Here, we carried out PheWAS in a Korean population using comprehensive health check-up data linked with genotype data, and furthermore identified applicable interpretations of the PheWAS results. We also compared the results of PheWAS studies conducted in different populations to evaluate trans-ethnic differences. Finally, our bipartite phenotype network analysis of phenotypes using shared genetic association revealed hidden patterns between phenotypes.

The deep phenotypes we used in our studies were corroborated during comprehensive health check-up by various confirmatory methods such as laboratory tests, endoscopy, CT scans, MRI, interview questionnaires, and so on. For each participant, all tests were done in the same institute and on the same day. This process of generating deep phenotypes makes for data quality that is well controlled and consistent when compared to results from phenotypes based on ICD codes, which can be discrepant with actual clinical diagnoses due to biases in billing pattern ^71^. As we were able to use the raw data produced by the test, our analysis included a lot of endophenotypes. Endophenotype (intermediate phenotype) is a quantitative biological trait ^72^ that is reported to reliably reflect the function of the categorical biological system ^72,73^ and has reasonable heritability ^74^. As such, an endophenotype could be more closely related to the genetic basis and cause of a clinical trait than would be a broad clinical phenotype such as an ICD code ^75^.

We compared our PheWAS results with studies done in European (UK Biobank) and Japanese (Biobank Project Japan) populations and found several novel loci, replicated loci, replicated phenotype-phenotype pairs. We furthermore compared estimated heritability among the populations. Significant variants in the Korean population were partly replicated in both European and Japanese populations, though the replication rate was higher in the Japanese population. We also identified SNP-phenotype associations that were unique to the Korean population when compared to not only the European but also the Japanese population. Noticeably, in the comparison of significant variants associated with body mass index (BMI), the Korean population had novel unique variants (Figure S8) associated with *TERF2IP, ATRNL1*, and *BANF1*. The results from these trans-ethnic and trans-nationality comparisons seemingly emphasize the importance of considering genetic differences among ethnicities, and also race. Koreans are generally included in the East Asian population; however, study of the human Y-chromosome ^76^ suggests that compared to other populations from Asia, the Korean population has characteristics of a distinct, mostly endogamous ethnic group, and living in a confined peninsula area has preserved these monogenic nationality traits. In a study comparing genetic structure and divergence among Han Chinese, Japanese, and Korean populations those three East Asian populations were shown to have distinct genetic make-up and could be distinguished based on their genetic characteristics ^77^. In the meta-analysis of our population and the Japanese population, 72.5% of phenotypes had variants that were uniquely significant in the meta-analysis. Our study shows that the common and exclusive genetic associations of phenotypes should be taken into consideration when performing a population-based clinical study. Furthermore, meta-analysis of PheWAS studies in populations of the same ethnicity but different nationalities can discover uniquely significant variants.

In the calculation of heritability for each phenotype, we generated a heritability estimation model that incorporated all phenotypes having cross-phenotype association through principal component analysis (PCA) into the 1st principal component (1st PC). For spondylosis, exercise, liver hemangioma, and gastroesophageal reflux, this estimated heritability using the 1st PC had higher explanation than did the value estimated without incorporating cross-phenotype associations. Heritability is the contribution of genetic variants that explains phenotypic variation ^78^. There is a well-known gap between the contribution calculated from genome-wide association results and that from classical heritability methods, termed missing heritability ^79^. The speculative explanations for explaining are a lack of statistical power of the variant ^80^, epistatic interactions, gene-environment interactions, and structural variations ^81–83^. The results in our study suggest that incorporating cross-phenotype association information for each phenotype could help fill out the missingness in heritability.

When comparing estimated heritability among different populations, the heritability in the Korean population of biomarkers for coagulation function, such as PT and aPTT, showed similar trends with that of the Japanese population, but manifested relatively high heritability when compared to the UK population. This indicates that the contribution of genetic variants to variation in coagulation traits is affected by ethnical differences. Evaluating heritability difference by ethnicity will be important supportive information in the development of drugs as an aspect of precision medicine.

We also leveraged the cross-phenotype association results to provide a panoramic overview of the network connections among multiple phenotypes and genetic variants. Specifically, we generated a phenotype-genotype network focused on metabolic syndrome (Figure 3A). Metabolic syndrome is a cluster of metabolic abnormalities that are known to be associated with visceral adipose obesity ^84^. A large number of epidemiological studies have been conducted on metabolic syndrome because it is a crucial target for healthcare, imposing an increased risk of developing conditions such as cardiovascular disease ^84^, malignant disease ^85^, depression ^86^, and metabolic disease ^84^. Early diagnosis is important to prevent the negative consequences of metabolic and this may be done by modifying the lifestyle and risk factors.

The network we constructed provided a rationale for defining metabolic syndrome by phenotypes of TG, HDL, hypertension, diabetes, and WC, and for using the characteristics of metabolic syndrome to collectively integrate heterogeneous and complex disease status. The network included phenotypes of cardiovascular disease (coronary calcium score, cardiac ischemia, brain atherosclerosis, malignant disease (thyroid cancer, gastric cancer), and depression and metabolic disease (fatty liver, uric acid), which are known to be complications of metabolic syndrome. Other phenotypes in the network related to obesity, specifically visceral obsesity indicator and visceral fat amount; obesity is a well-known cause of metabolic syndrome ^84^. Furthermore, lifestyle factor phenotypes such as alcohol consumption, smoking habit, and exercise amount were also part of the network. These suggest modifiable targets for preventing the complications of metabolic syndrome. Finally, the network suggested hub genes associated with metabolic syndrome. Similar network analysis of PheWAS results might provide genotype-based evidence of connections among phenotypes or variants, which to date have been assumed from epidemiological research, and can also provide novel insights into connections that have not been previously reported or recognized.

We additionally used the bipartite phenotype network to perform cross-phenotype mapping. Table S23 shows the cross-phenotype mapping constructed for tumor markers. Tumor markers are highly used in clinical practice for tasks such as oncological screening and monitoring recurrence after treatment. The marker carcinoembryonic antigen (CEA) is recommended by the National Comprehensive Cancer Network (NCCN) guidelines for colon cancer and American Society of Clinical Oncology (ASCO) to test a diagnosis of colon cancer as a baseline for monitoring and then to regularly monitor for recurrence or metastasis of the colon cancer ^87,88^. Testing for the marker PSA is recommend by the American Cancer Society (ACS) for men aged >50 years, after an informed decision-making process ^89^. Regular testing for another marker, serum alpha-fetoprotein (AFP), is recommended by the NCCN guideline in the follow-up of hepatocellular carcinoma ^90^.

However, while testing for tumor markers is essential in the surveillance of malignant disease, their usage faces problems in the form of low sensitivity and specificity and the potential that they could be affected by factors other than the cancer itself. Thus, providing a cross-phenotype mapping for tumor markers could support an oncologist in interpreting the results of each tumor marker test. For instance, hemoglobin was included in our CEA cross-phenotype mapping. Thus, if a colorectal cancer patient has severe anemia, we should be cautious about interpreting a change in CEA; the anemia could attenuate or exaggerate its reflection of the patient’s cancer status ^60^. There are several reports that have used not only one tumor marker but a combination of tumor markers to monitor malignancies ^91–93^. In Table S23, each tumor marker has pairs with multiple other tumor markers, which provide supporting evidence for combining tumor markers as a means to improve their utility in malignancy surveillance.

We also built cross-phenotype mappings for environmental factors. Figure 3C shows the cross-phenotype mapping for coffee consumption in particular. Similar visualization of the correlations between environmental factors and other phenotypes could provide insight into which disease should be considered for the investigation of the benefits or hazards of given environmental factors, and what also connections could provide a candidate model for gene x environment interactions.

In our study, we applied Mendelian randomization analysis to cross-phenotype networks in order to generate corresponding causal inference networks. To the best of our knowledge, this is the first approach to utilize MR in network-based analysis. MR enables the estimation of causal inference by evaluating the relationship between genetic susceptibility to the causal factor and the outcome in question ^94^. As shown in Figure 3D, we specifically drew a causal inference map for skeletal muscle mass. We visualized this map because skeletal muscle mass is regarded as an endocrine and paracrine organ, and is also suggested as a marker in diseases such as metabolic syndrome, diabetes, and more ^95^. The analysis revealed skeletal muscle mass as having significant causal inference for metabolic syndrome. Thus, by performing MR, we can suggest which phenotype could be causal or an outcome in relation with a trait and also begin to elucidate the mechanism or pathophysiology for a disease of interest.

Our study has several advantages. First, to the best of our knowledge, this is the first PheWAS study performed in the Korean population. As described above, several loci in this population differ from the Japanese population. We were also able to carry out trans-nationality analysis for the PheWAS. Second, we defined phenotypes directly using results from health check-ups and questionnaire responses from personal participants. This make the resolution, clarity, and reliability of this study’s results better than those of a PheWAS based on ICD codes. Third, since all tests were performed in the same institute, under the same conditions, and by using the same machines, protocols, and chemicals, the produced data is consistent and its quality is highly controlled. Fourth, we tried to translate the PheWAS results in ways meaningful to healthcare practice and clinical research, so that the results could be highly applicable and utilized more practically. We constructed a phenotype-phenotype network using all the phenotypes in our study (Figure S13). Similarly constructing a phenotype-phenotype network based on comprehensive, deep phenotypes could provide clinicians and researchers with a detailed landscape of the interconnections between phenotypes and enable better understanding of their underpinnings. Furthermore, the phenotype-phenotype network not only includes disease status but also contains information on genes, environment, and lifestyle. Precision medicine pursues prevention and treatment strategies that take individual variability ^1^, such as in genes, environment, and lifestyle, into account ^2^. Accordingly, the networks generated by PheWAS would provide fundamental information for realizing precision medicine in healthcare practice and clinical research.

Our study has several limitations. First, we did not have a set Korean replication population because it was not possible to find such datasets with the variety of deep phenotypes incorporated in our study. However, we instead introduced the UKBB and BBJ as replication sets, and consequently identified multiple replicated loci. We also replicated the phenotype-phenotype pairs using a larger EHR-driven database of Korean samples to investigate whether the genetic connection was reflected at the actual phenotype level. Second, the study population was collected from those who had regular health check-ups, and therefore samples with positive morbidity were relatively few. Accordingly, the significance of the loci was low for some phenotypes. We tried to overcome this lack of statistical power by performing a meta-analysis with the UKBB and BBJ summary statistics, in which we were able to pick up additional significant loci. In a future study, we will incorporate diverse disease cohorts from the Korean population to increase the study power.

In conclusion, our study highlights the capacity for realizing interpretable clinical applications of PheWAS through comprehensively exploiting the results. With the information generated by PheWAS, we attempted to provide a landscape that integrated an individual’s genetic, lifestyle, and environmental factors along with health status. We provided several samples of actionable applications such as constructing a gene-phenotype association network related to metabolic syndrome; constructing cross-phenotype mappings; and visualizing causal inference mappings. Through analysis in the context of differences in ethnicity and nationality, our study shows that some phenotypes are common or exclusive in their genetic associations, and this should be taken into consideration when performing a population-based clinical study.

The paradigm of PheWAS suggested in our study will eventually be the cornerstone for applying the core concepts of precision medicine to research and healthcare practice.

## Supporting information

Index to supplementary materials

## Acknowledgement

We acknowledge Jong-Eun Lee of DNA Link for the collaborative support to establish the GENIE Cohort.

## Data availability

Complete summary statistics are not publicly available due to restrictions (institutional policy to protect the privacy of research participants), but are available from the corresponding author on reasonable request. However, all other data are contained in the article and its supplementary information or are available upon reasonable request.

