## Supplementary material for "Phenome-wide association study of a comprehensive health check-up database in a Korea population: Clinical application & trans-ethnic comparison": Index to supplementary materials

**<Supplementary Figures>**

**Figure S1.** Imputation Performance evaluation by NARD+1KG reference platform in SNPs with MAF>0.01

**Figure S2.** Principal component analysis for population stratification

**Figure S3.** Pipeline of the data preprocessing in the study

**Figure S4.** Systematic analysis of the PheWAS results

**Figure S5.** A phenome-wide Manhattan plot of PheWAS for associations with P value less than 10^-4^ in all phenotypes

**Figure S6.** Functional annotation (significant loci, p value < 1x10^-4^)

**Figure S7.** The schematic structure of the trans-ethnical

**Figure S8.** Trans-ethnic, trans-nationality comparison of PHEWAS

**Figure S9.** Trans-ethnic, trans-nationality comparison of PHEWAS : Comparing the significant loci and gene for Body mass index (BMI), among Korean, Japan, UK population

**Figure S10.** Trans-ethnic, trans-nationality comparison of PHEWAS : Comparing the phenotype-phenotype pairs in bipartite phenotype network

**Figure S11.** Applicable interpretation - Estimated heritability

**Figure S12.** Applicable interpretation - Comparing the heritability among Korean, Japan, UK population

**Figure S13.** Applicable interpretation - Network analysis

**Figure S14.** Applicable interpretation - Relationships among obesity indices

**Figure S15.** Applicable interpretation - Cross phenotype mapping

**Figure S16.** Meta-analysis of phenome-wide associations studies in Korean and Japanese population

**Figure S17.** Manhattan plot for the unique significant variants from meta-analysis

**<Supplementary Tables>**

**Table S1.** Glossary for the phenotypes used in analysis

**Table S2.** Comparing the imputation accuracies using different reference panel in chromosome 22

**Table S3.** Matching common phenotypes among population

**Table S4.** Characteristics of the subjects enrolled in this study

**Table S5.** Phenome wide association study results with number of significantly associated loci and genes

**Table S6**. Top significant loci in PheWAS, for discrete and continuous phenotype

**Table S7.** Top 5 significant loci in PheWAS, for each phenotype

**Table S8.** functional annotation for the significant loci with p <10-4

**Table S9.** Details of the loci with high impact biological consequences

**Table S10.** Comparison of the PheWAS result from GENIE cohort (Korean) and BBJ (Japanese) in gene level / Comparison of the PheWAS result from GENIE cohort (Korean) and UKBB (European) in gene level

**Table S11.** Comparison of the PheWAS result from GENIE cohort (Korean) and BBJ (Japanese) in loci level / Comparison of the PheWAS result from GENIE cohort (Korean) and UKBB (European) in loci level

**Table S12.** Comparison of the PheWAS result from GENIE cohort (Korean) and BBJ (Japanese) in gene level significantly associated with body mass index

**Table S13.** Genes associated with body mass index, which are exclusive in Korean population comparing both Japanese and European population

**Table S14.** Possible polygenicity

**Table S15.** Possible pleiotropy

**Table S16.** Bipartite phenotype networks

**Table S17.** Summary of the degree properties for core phenotype in bipartite phenotype network

**Table S18.** Summary of the degree properties for core gene in bipartite gene network

**Table S19.** Pairs of phenotypes from bipartite phenotype network observed in Korean, Japanese and European population

**Table S20.** Summary of the heritability analysis

**Table S21.** Mean heritabilities of phenotypes in category

**Table S22.** Relationship among obesity indices based on cross phenotype associations (Refer Figure 3B and Figure S14)

**Table S23.** Cross phenotype mapping for tumor markers

**Table S24.** Significant mendelian randomization results for phenotypes in bipartite phenotype network (FDR<0.05 shown)

**Table S25.** Mendelian randomization analysis for life style factors as exposure (FDR<0.05 are shown)

**Table S26.** Correlation analysis in EHR-based HPEACE cohort

**Table S27.** Comparison of the phenotype-phenotype pairs between PHEWAS driven vs. EHR-driven

**Table S28.** Meta-analysis for PheWAS from Korean, Japanese and European population
