## Supplementary_Figures for "Phenome-wide association study of a comprehensive health check-up database in a Korea population: Clinical application & trans-ethnic comparison"

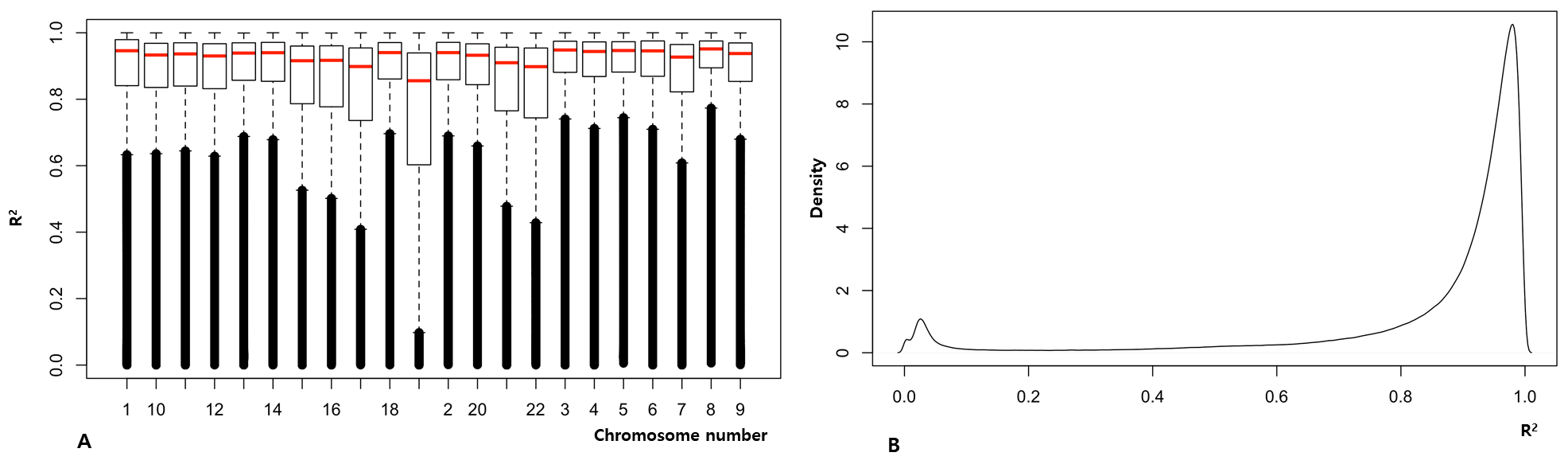

**Figure S1. Imputation Performance evaluation by NARD+1KG reference platform in SNPs with MAF>0.01**

(A) Box plot shows R^2^ distribution, which are calculated by the true genotypes and the imputed ones, for each chromosome

(B) Density plot shows density of R^2^ in all chromosomes

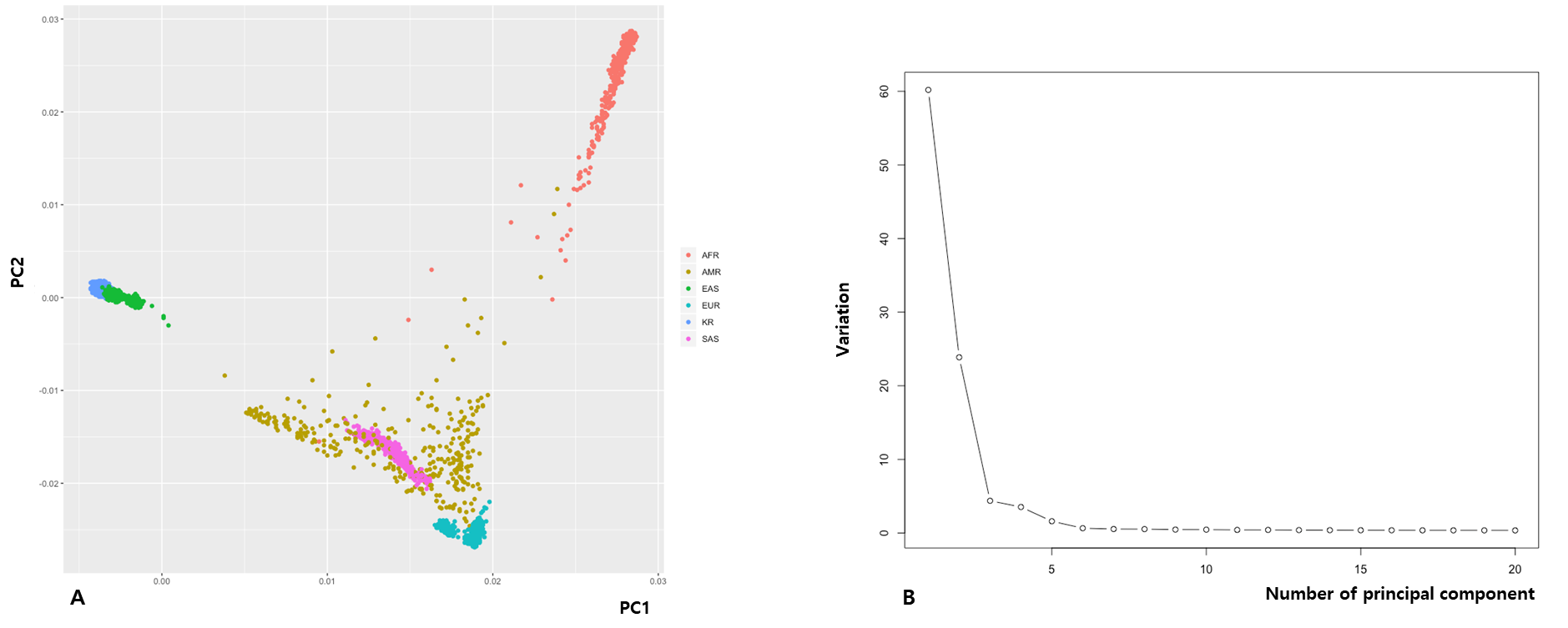

**Figure S2. Principal component analysis for population stratification**

Analysis of population stratification was performed to assess the influence of ethnicity (A) Principal component analysis (PCA) of global populations from AFR, AMR, EAS, EUR and SAS stand for Africans, Americans, East Asians, Europeans, and South Asians, respectively. The population of our study is assumed the representative of Korean population (KR). The total population was merged with AFR, AMR, EAS, EUR, and SAS data from the 1000 Genomes data for PCA and the KR showed distinct clustering. (B) We used the top three principal components (PCs) to adjust the population stratification (B)

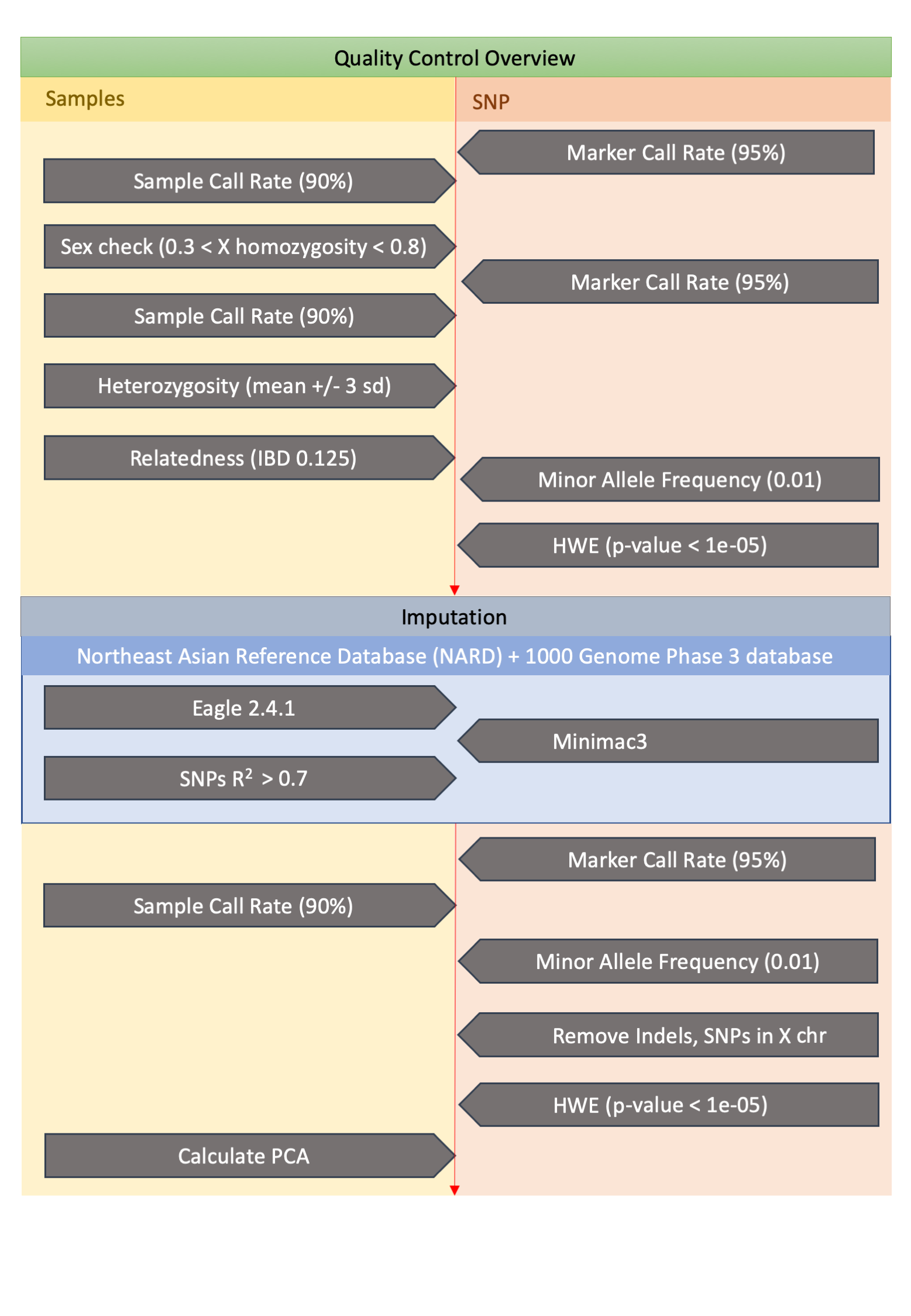

**Figure S3. Pipeline of the data preprocessing in the study**

We performed a quality control process (upper box) for sample (left box) and marker (right box). Imputation was done using the northeast Asian reference database and 1000 genome phase 3 database (lower box). After the process, 9,742 samples and 6,860,342 SNPs were used for the analysis

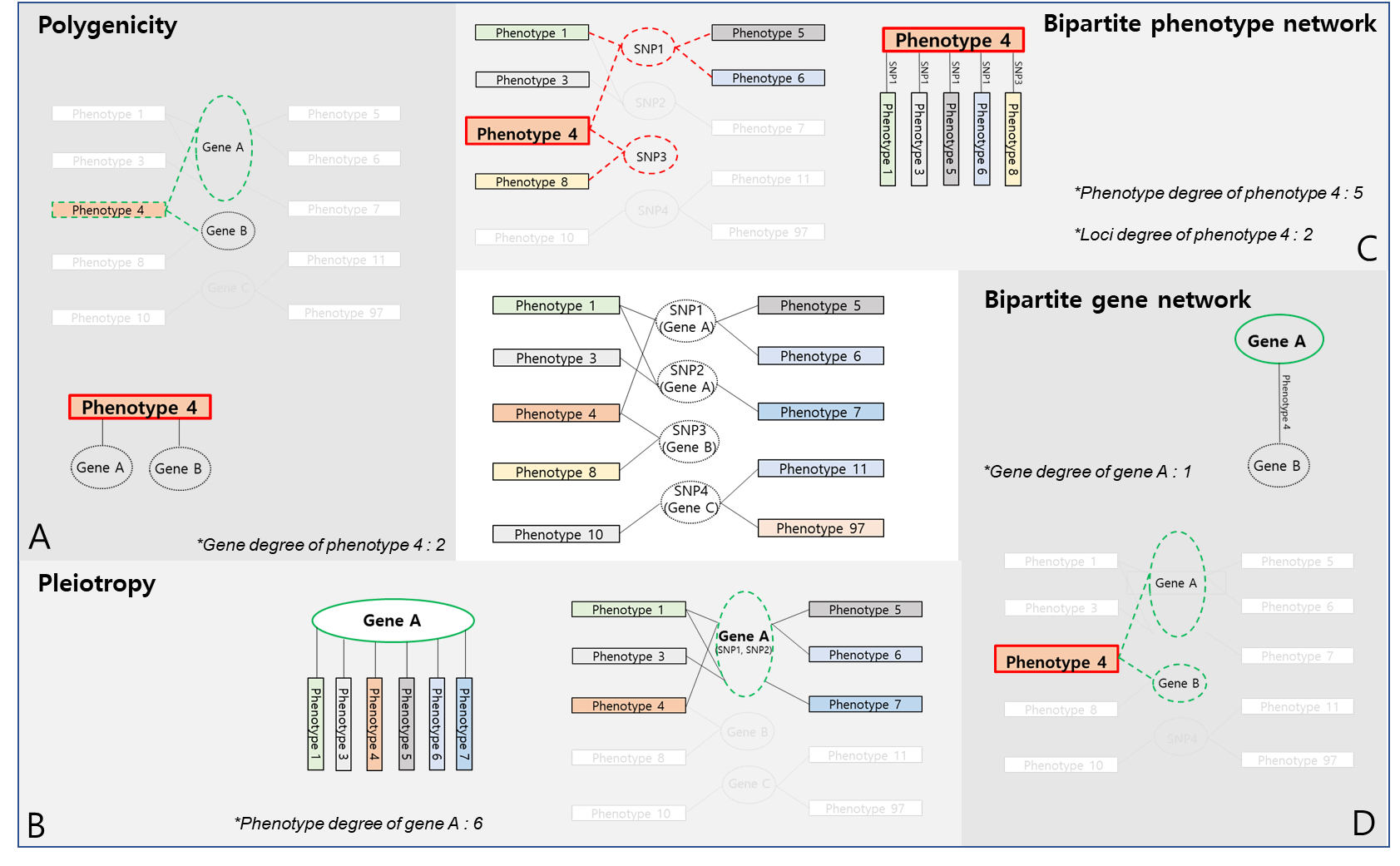

**Figure S4. Systematic analysis of the PheWAS results**

We leveraged the cross- phenotype associations, in which one loci is associated with multiple phenotypes, to perform systematic analysis of the PheWAS results

A. Polygenicity is one whose phenotype is influenced by more than one gene. In our study, the possible polygenicity is suggested. In the figure, Phenotype 4 is associated with Gene A and Gene B.
B. Pleiotropy is a definition in which a locus or a gene affects more than one phenotype. In our study, the possible pleiotropy is suggested. In the figure Gene A or SNP1 is association with multiple phenotypes.
C. Bipartite phenotype network was constructed by the connection among phenotypes sharing at least one loci. In the figure, Phenotype 4 has connection with five phenotypes (Phenotype 1,3,5,6,8) via the sharing associations with SNP 1 and 3
D. We defined bipartite gene network as the connection among genes shared by at least one phenotype. In the figure phenotype 4 is shared by both Gene A and Gene B.

In the connections or network, the degree property is the number of direct connections between one component and another component. In Figure S4C, Phenotype 4 has five phenotypes connected, that would result in 5 degree of phenotype for Phenotype 4. Phenotype 4 has two loci (SNP1 and SNP3) associated, which will mean 2 loci degree of Phenotype 4. In Figure 4SD, Gene A has one gene in connection, which will result one degree of gene for gene A.

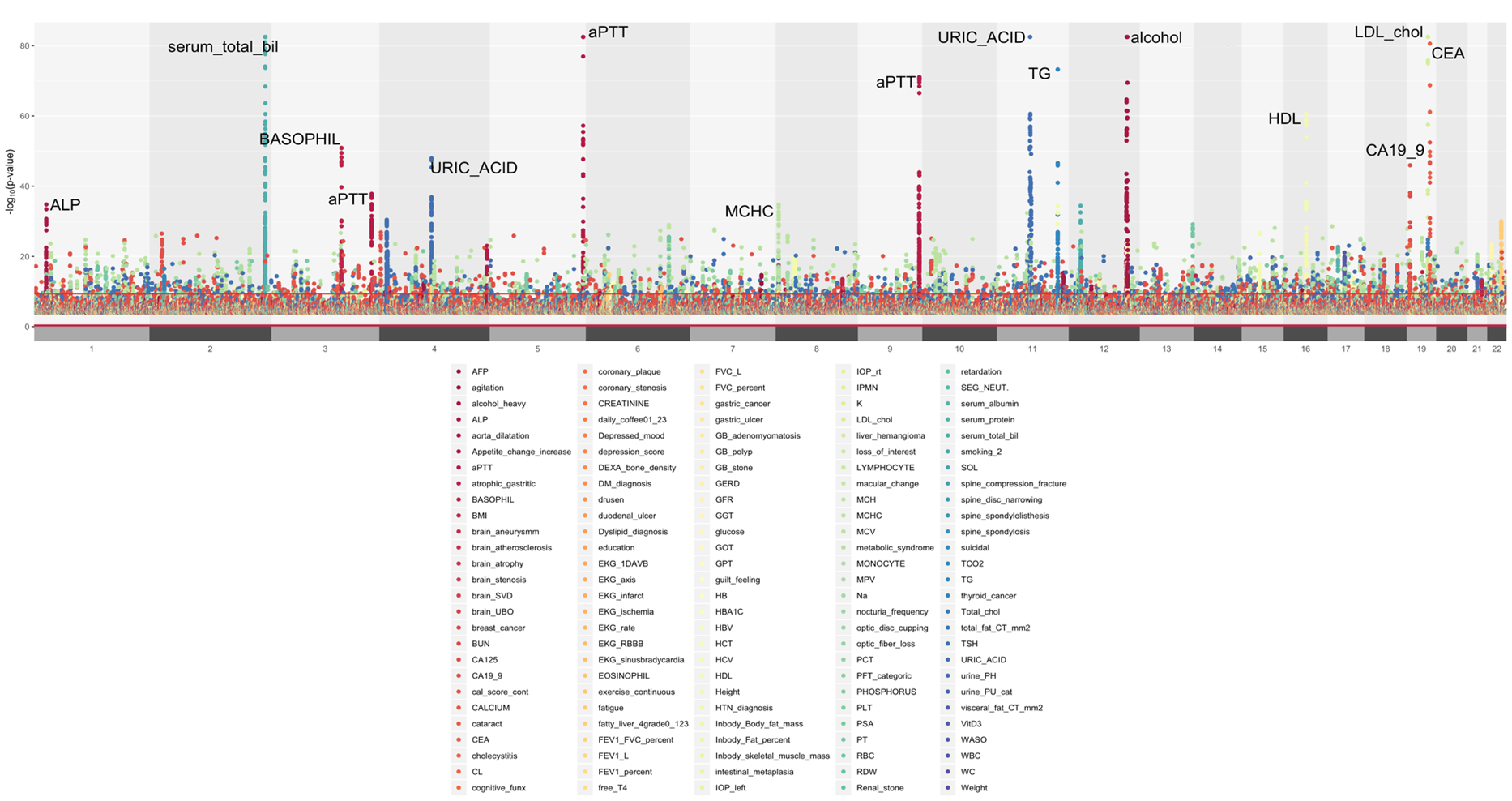

**Figure S5. A phenome-wide Manhattan plot of PheWAS for associations with P value less than 10-4 in all phenotypes.** The phenotypes with top significant p value associated loci were alcohol consumption (Alcohol), activated Partial Thromboplastin Time (aPTT), carcinoembryonic antigen (CEA), LDL cholesterol (LDL), serum total bilirubin (Tbil), uric acid (Urate), triglycerides. aPTT had two spots of top signal loci in the plot, in chromosome 5 and chromosome 9. Each color of the dots denotes the respective phenotypes.

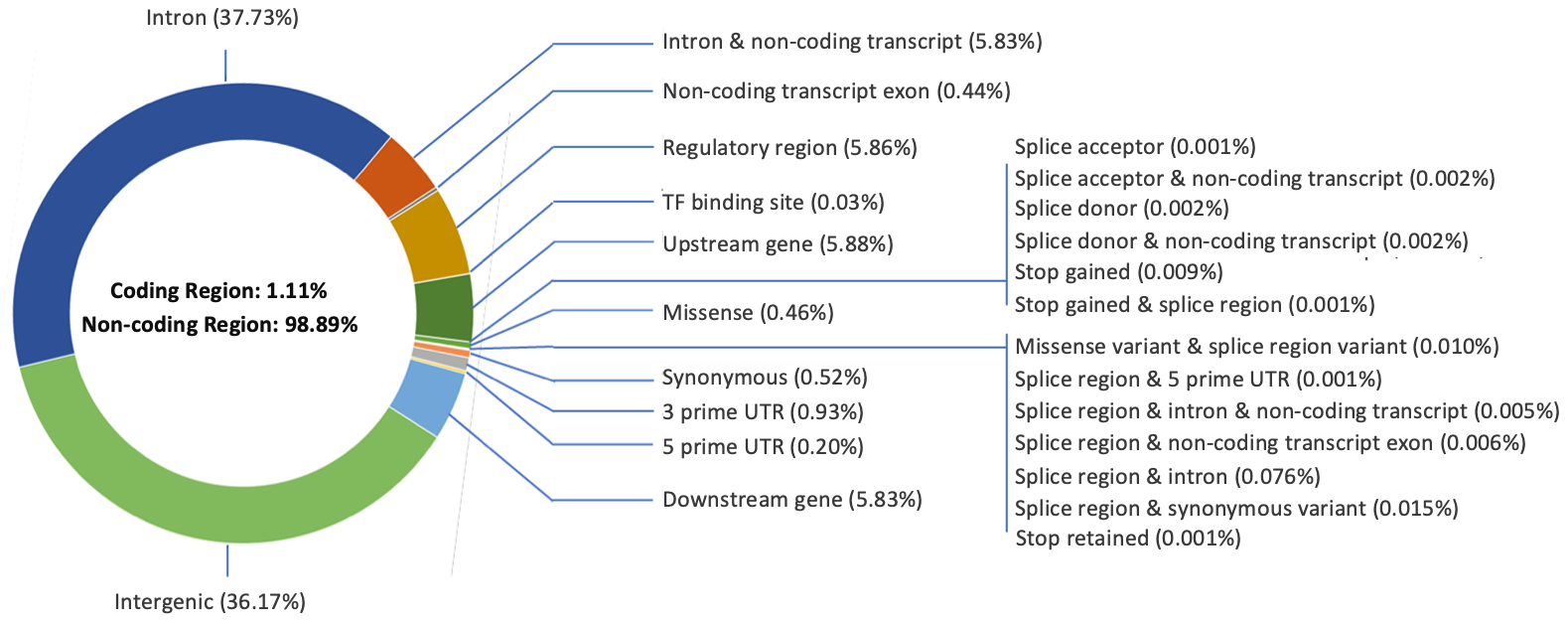

**Figure S6. Functional annotation (significant loci, p value < 1x10^-4^)** We annotated the functional consequences of the loci on genes or proteins provide by Ensemble variant effect predictor (VEP). The plot shows the number of variants in each type of predicted consequences for the loci with p value less than 1x10-4 in PheWAS.

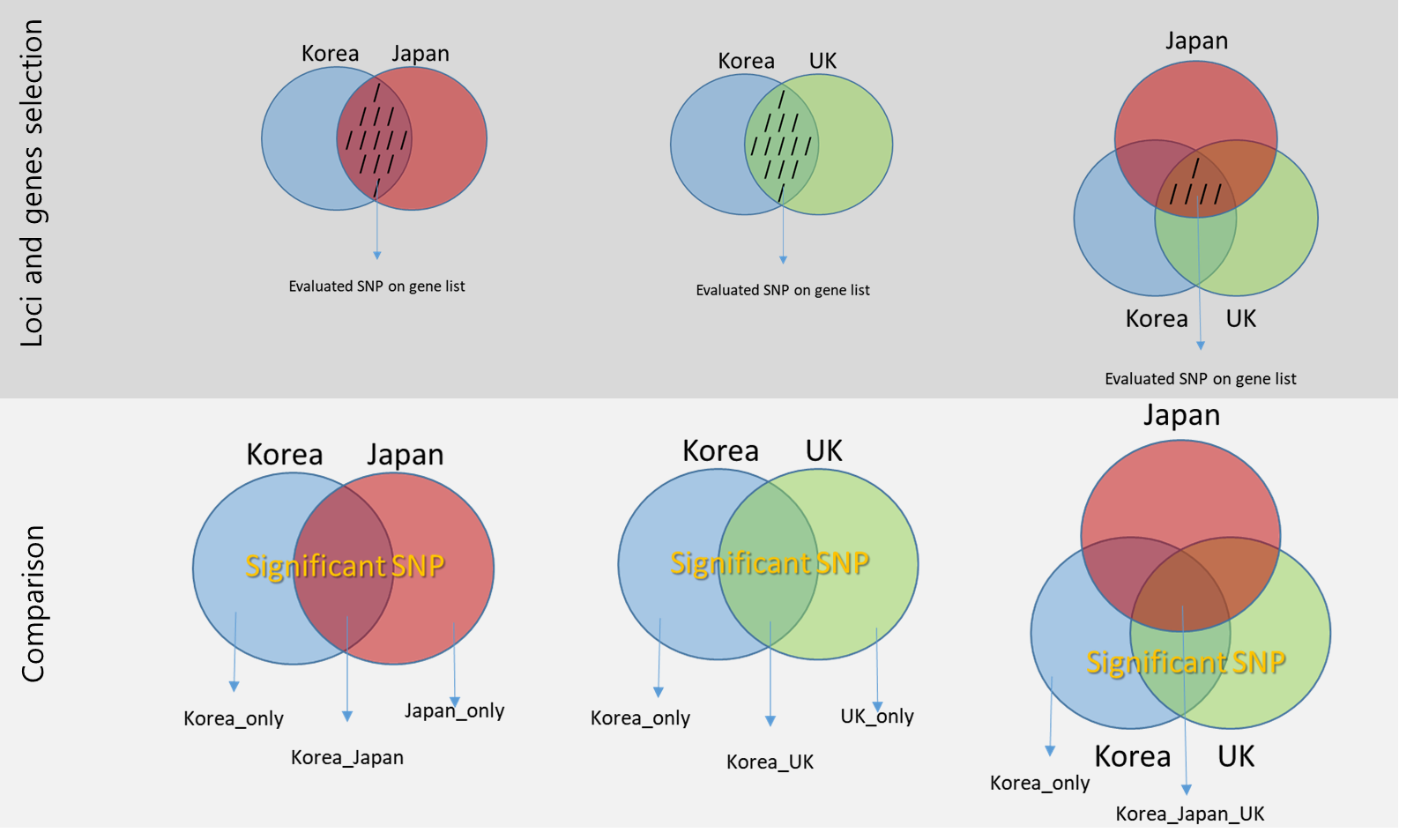

**Figure S7. The schematic structure of the trans-ethnical**

We utilized the results from UK biobank and BioBank Japan Project to perform trans-ethnic, and trans-nationality comparison of the PheWAS results **A.** Since each population used different SNP array platform, the loci used in the analysis were different. We filtered the loci used in both of the populations. In the figure, the comb marked area is the loci or genes used for comparison. **B.** We compared the significant loci (p <10-4) among the populations. The overlapping sector between our population and others could be assumed as the replicated loci of our PheWAS results. Korea only sector could be the novel loci or gene found in our PheWAS results. The overlapping sector will be the replicated loci or genes.

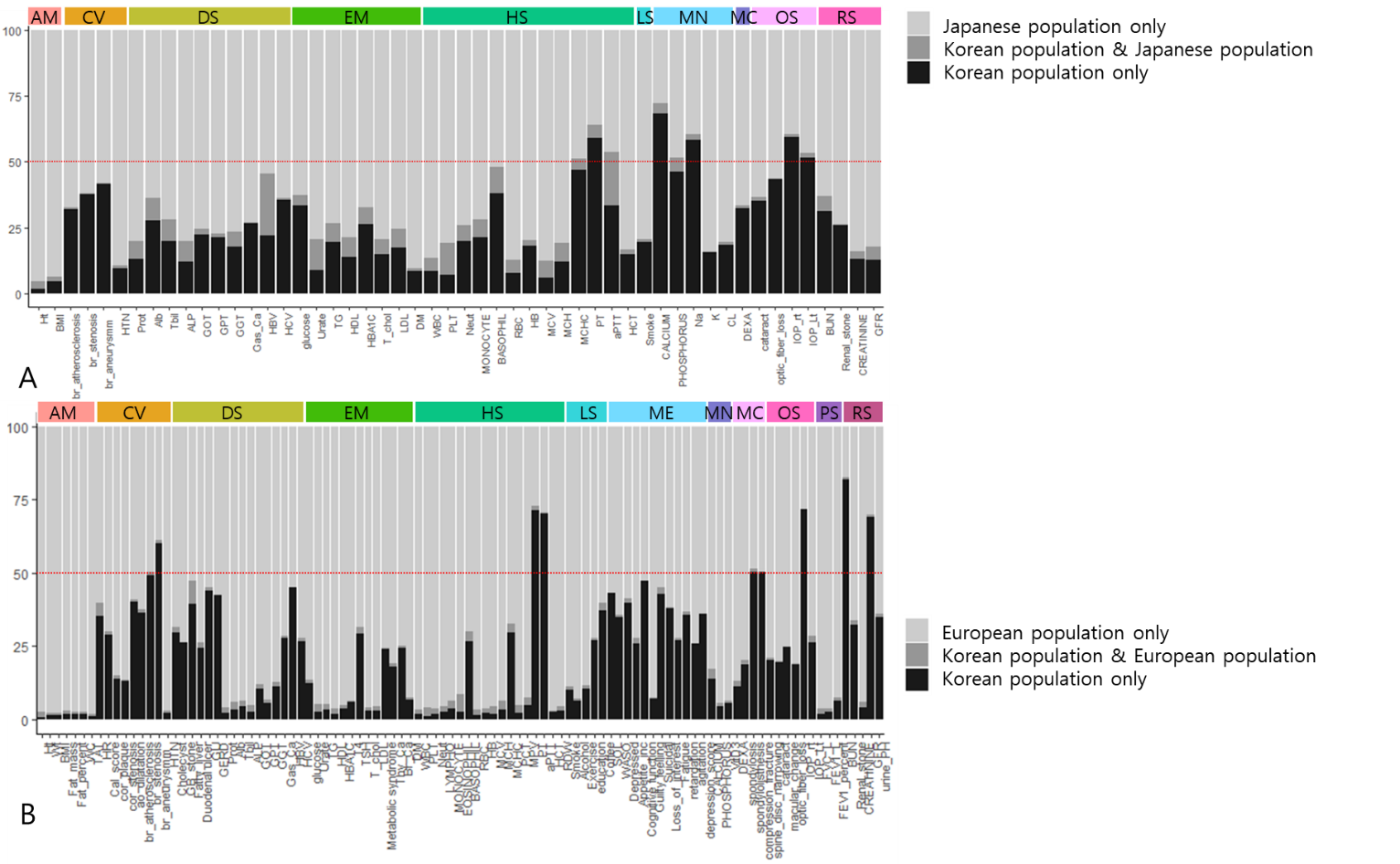

**Figure S8. Trans-ethnic, trans-nationality comparison of PHEWAS**

We compared the PHEWAS results among Korea, Japanese and European. The phenotypes existing in both populations were used. We evaluated the genes significantly associated only in Korean (Black bar) vs in both population (gray bar) vs. only in Other population (bright gray bar). The colored bar at the top is the category for the phenotypes. The Y axis denotes the ratio (%) of each classification of loci, putting 100% as total loci significant in Korea population or other population. **A.** PHEWAS result comparison between Korean population vs. Japanese population. **B.** PHEWAS result comparison between Korean population vs. European population.

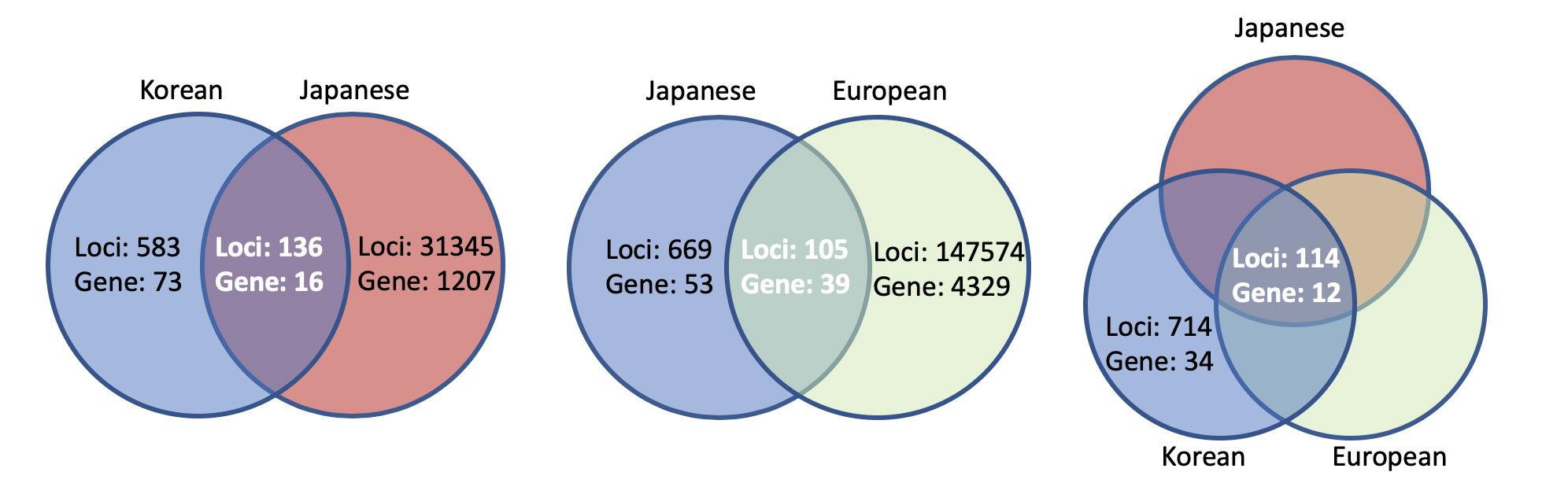

**Figure S9. Trans-ethnic, trans-nationality comparison of PHEWAS : Comparing the significant loci and gene for Body mass index (BMI), among Korean, Japan, UK population**

In the association study for body mass index (BMI), 136 loci (16 genes) were replicated in Japanese population and 105 loci (39 genes) in European population, respectively. There were 34 genes (714 loci) unique in Korean population comparing both Japanese and European population.

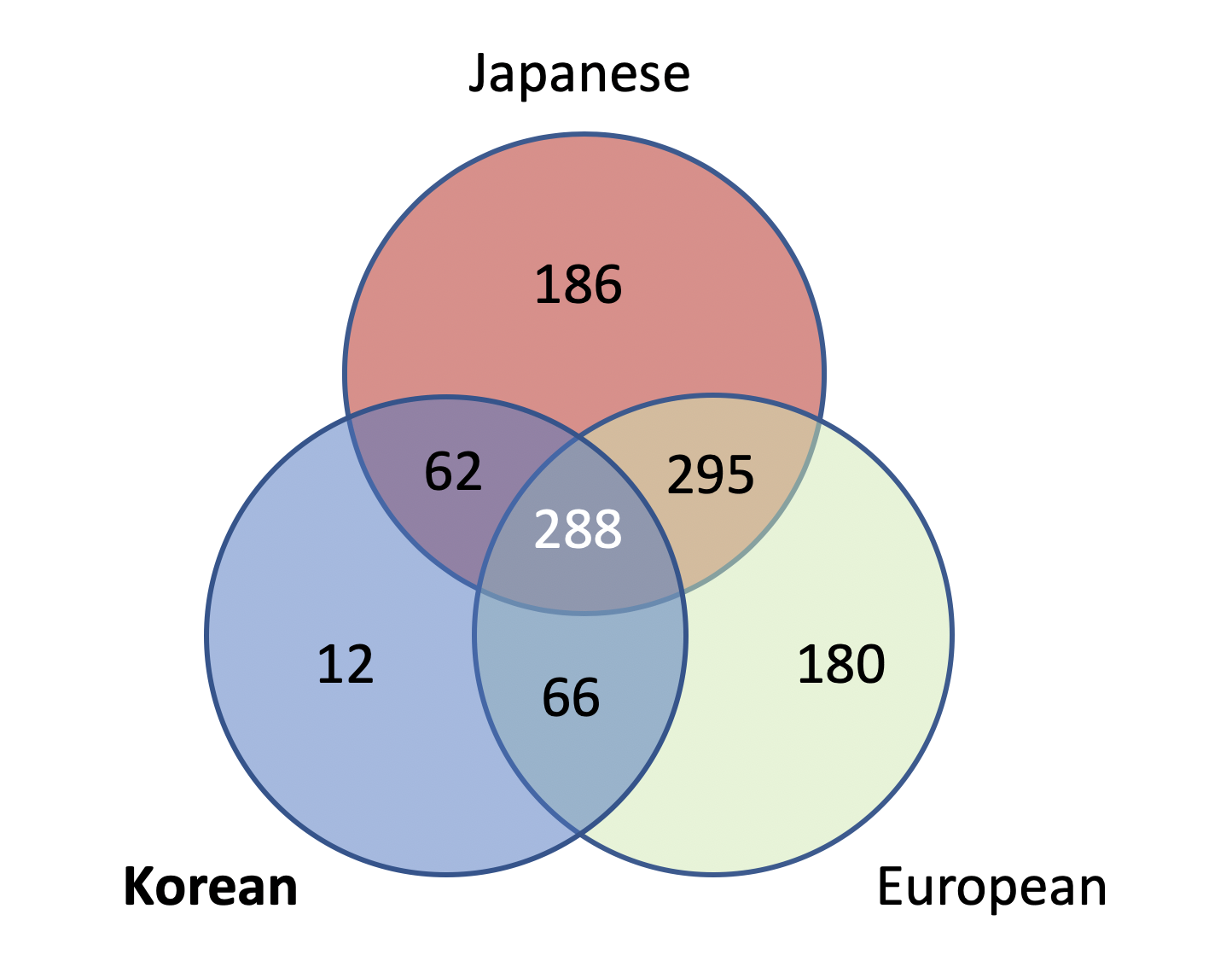

**Figure S10. Trans-ethnic, trans-nationality comparison of PHEWAS : Comparing the phenotype-phenotype pairs in bipartite phenotype network**

There are 49 phenotypes that are analyzed in PheWAS for all population. The bipartite phenotype network was generated by these phenotypes. In the venn-diagram, there were 288 pairs of phenotypes that were simultaneously observed in all three populations.

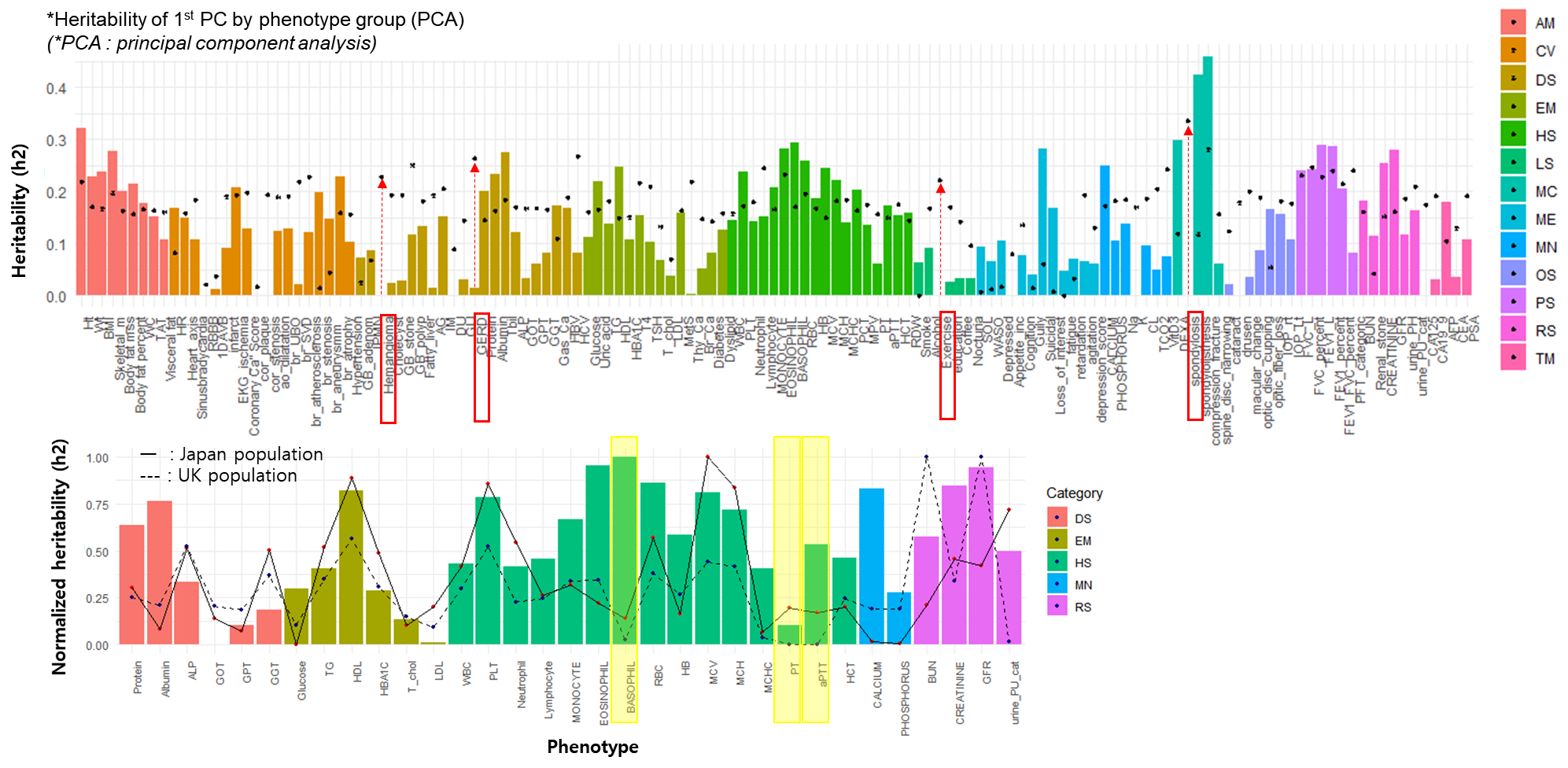

**Figure S11. Applicable interpretation - Estimated heritability**

A. The heritability for each phenotype was calculated using LD score regression, which is shown as bar plots. The colors correspond to the category of the phenotypes included. As a further evaluation, we performed a principal component analysis (PCA) for each phenotype using the phenotypes connected by bipartite phenotype network. We used the 1^st^ principal component (1^st^ PC) to calculate the heritability for each phenotype, which is shown as *. Spondylosis, exercise, liver hemangioma, gastroesophageal reflux, and etc. had improvement of heritability by extracting the features generated by the phenotypes in bipartite phenotype network. **B.** We compared the heritability of the phenotypes observed simultaneously in GENIE, UKBB and BBJ to see the difference of heritability in Korea, European and Japanese. Since each population used different SNP array platform and loci to calculate heritability, we normalized the heritability and compared the trends in heritability. The bar plot is the heritability for Korea and the straight line for Japanese and dashed line for European. In Korean, heritability of basophil (%) was relatively high compared to both populations. The phenotypes related to coagulation function, PT and aPTT, were relatively high compared to UKBB population but similar to BBJ population.

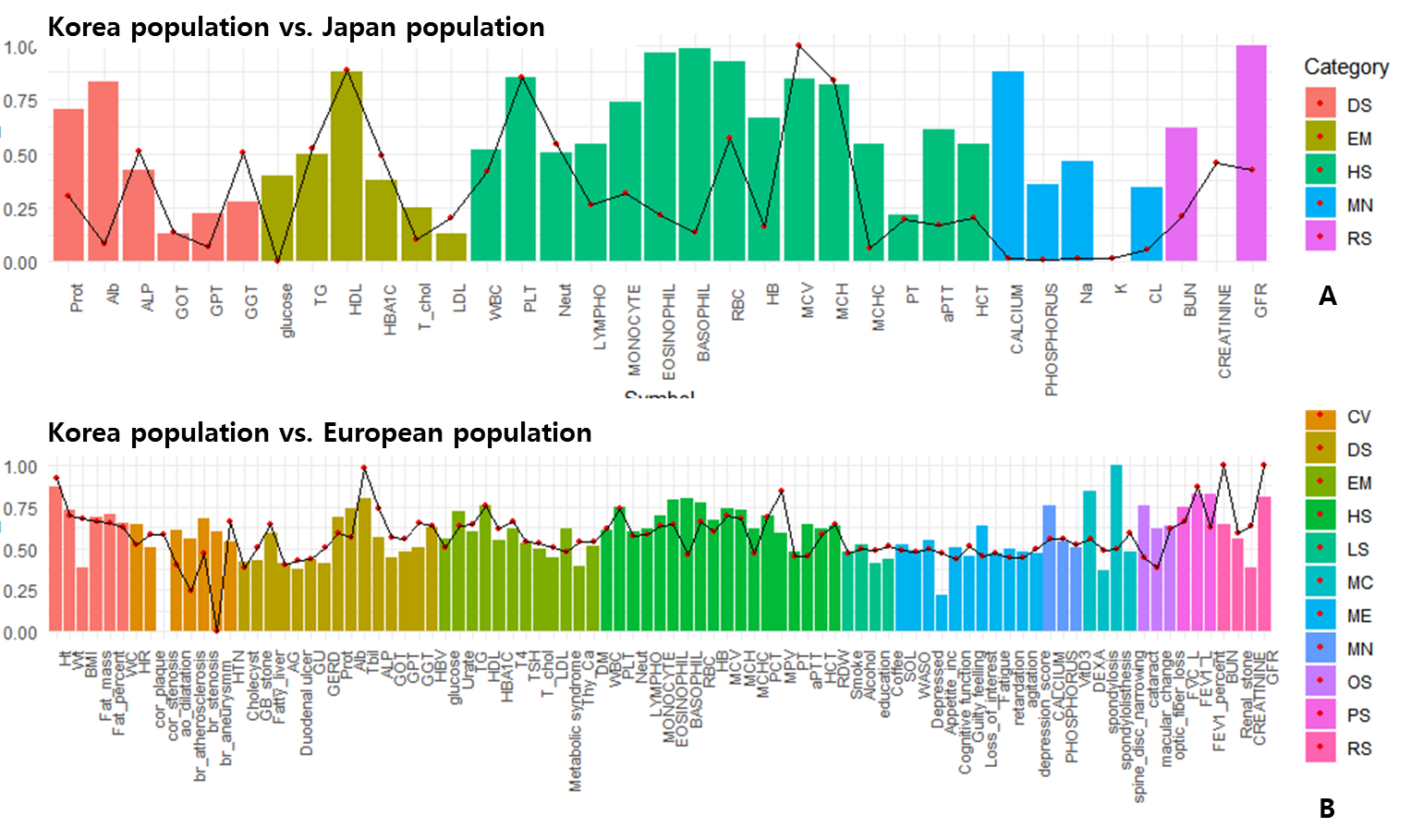

**Figure S12. Applicable interpretation - Comparing the heritability among Korean, Japan, UK population**

**A.** The calculated heritability between Korean population (bar plot) and Japanese population (line plot) is shown. **B.** The calculated heritability between Korean population (bar plot) and European population (line plot) is shown.

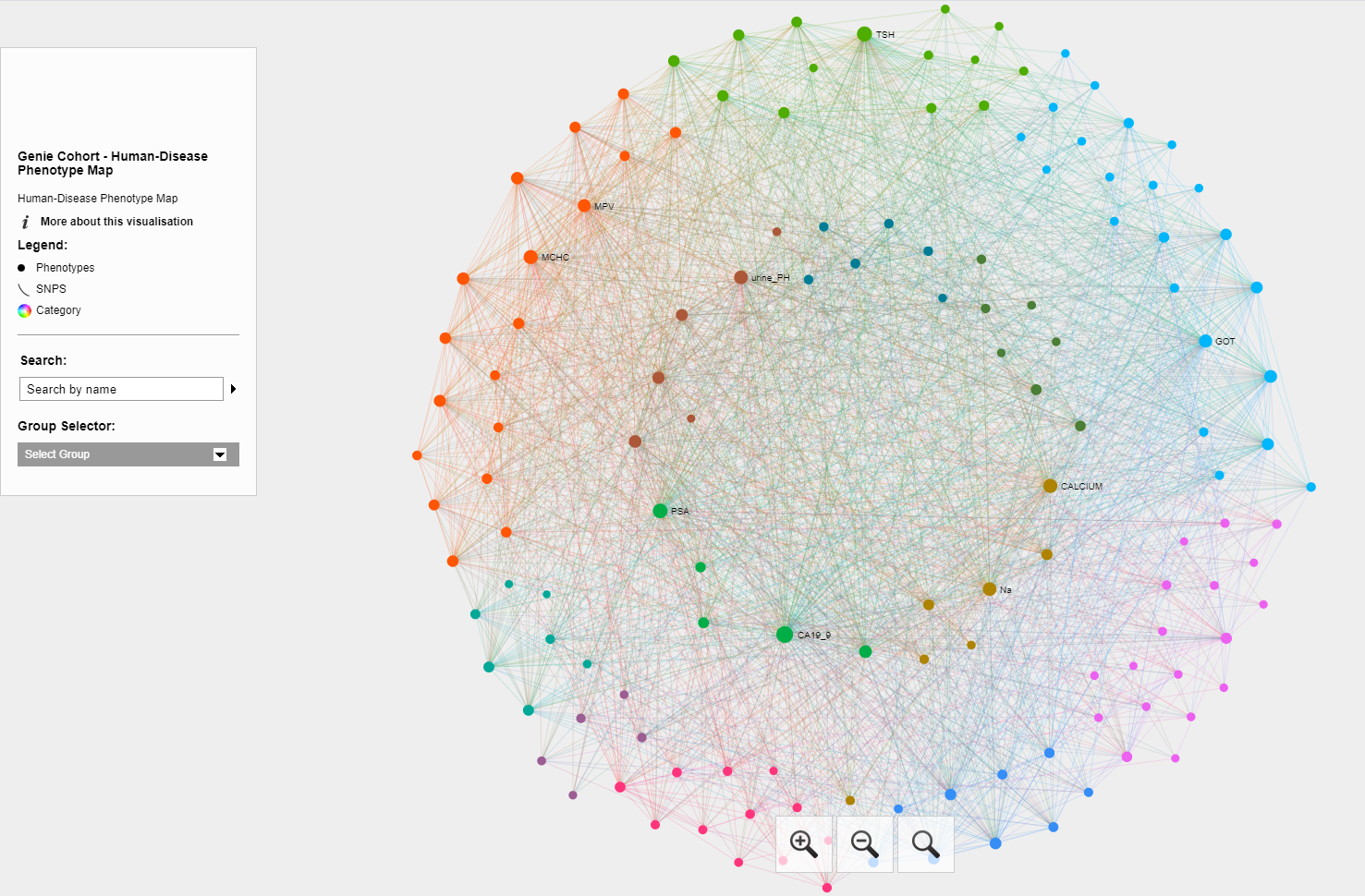

**Figure S13. Applicable interpretation - Network analysis**

Using the 1,926 pairs of phenotypes, based on sharing same loci (P<1x10^-4^), we constructed a phenotype-phenotype network for our dataset.

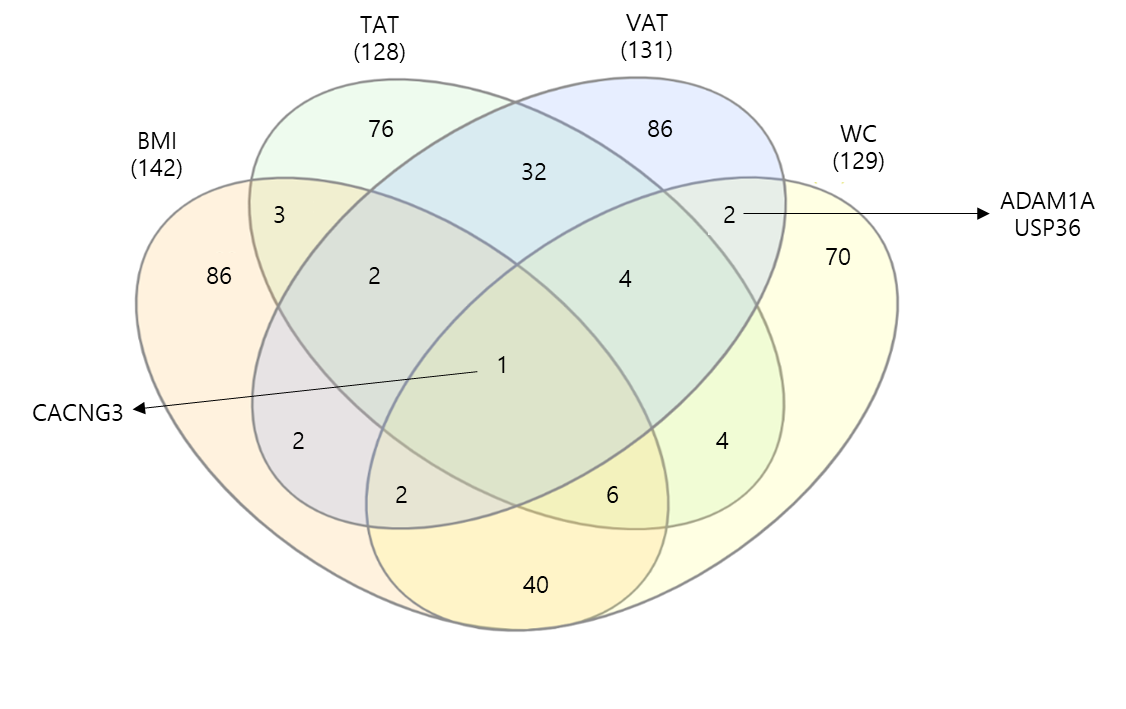

**Figure S14. Applicable interpretation - Relationships among obesity indices** We visualized the comparison among the obesity indices such as body mass index (BMI), waist circumference (WC), visceral adipose tissue (VAT) and total adipose tissue (TAT) amount by drawing a the venn-diagram for cross phenotype association of genes. 14 phenotypes were associated exclusively with VAT and WC and this intersection had 2 genes associated.

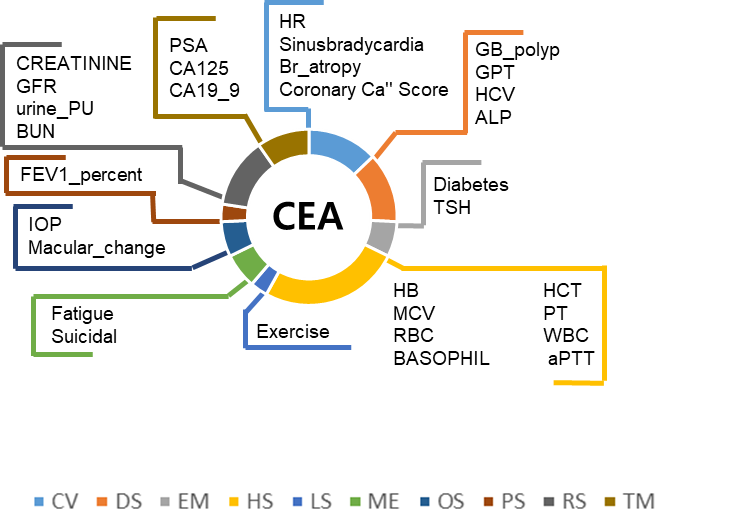

**Figure S15. Applicable interpretation - Cross phenotype mapping** Cross phenotype mapping was generated base on the bipartite phenotype network, constructed by the connection among phenotypes sharing at least one loci. Tumor markers, CEA, CA19-9, AFP, PSA, CA125, had high degree of phenotype in the bipartite phenotype network. The figure shows the cross-phenotype mapping for tumor marker CEA, which could be considered during the oncological practice and taking consideration of all the possible effects of phenotypes other than colorectal cancer progression itself.

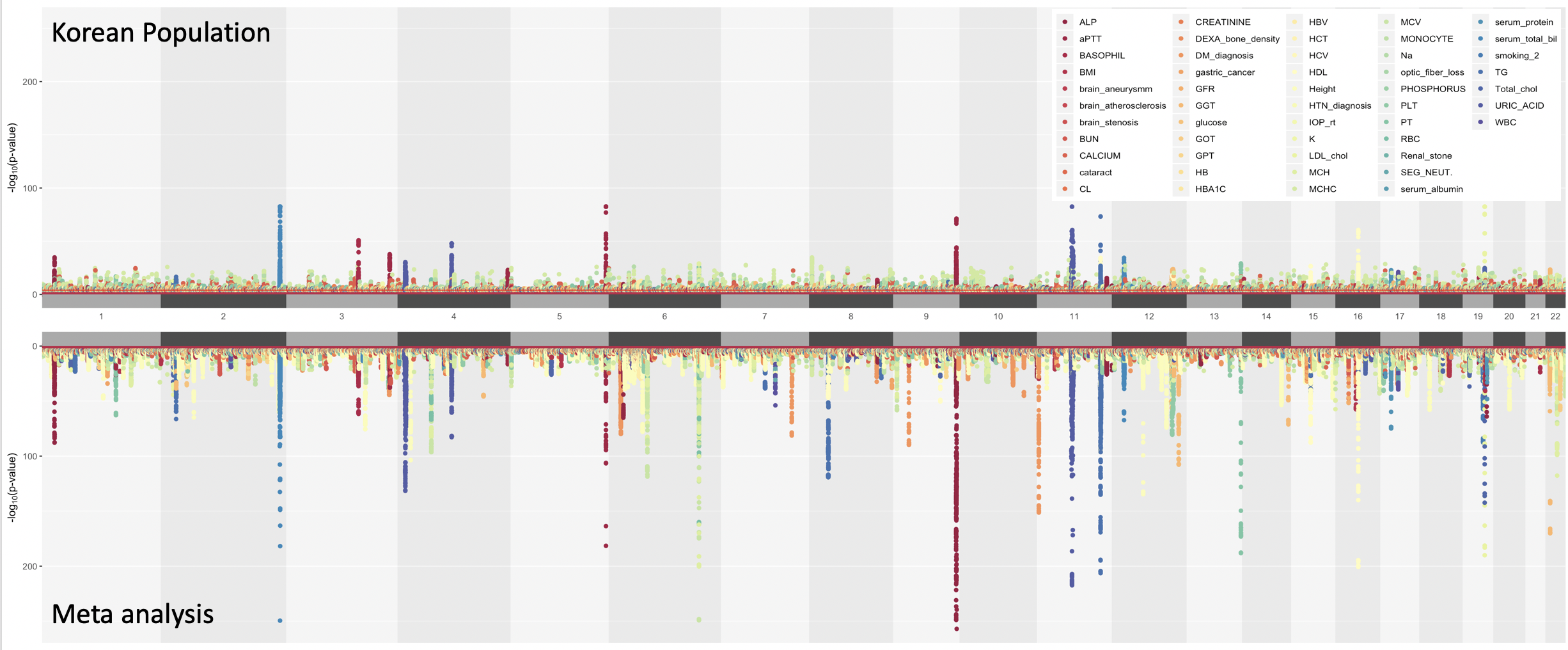

**Figure S16. Meta-analysis of phenome-wide associations studies in Korean and Japanese population** The upper plot is the PheWAS results done in Korea population and the lower plot is the meta-analysis result by METAL

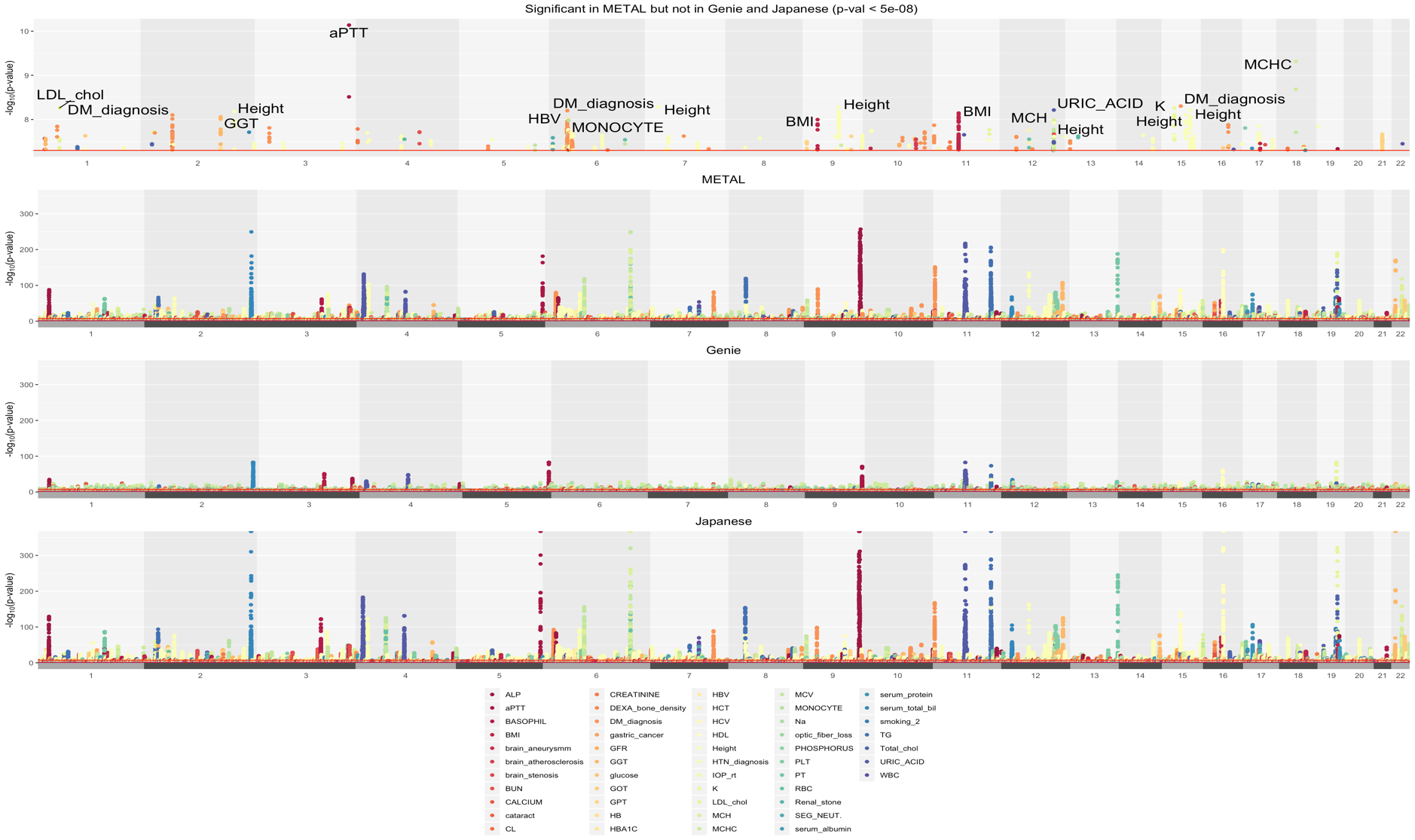

**Figure S17. Manhattan plot for the unique significant variants from meta-analysis :** The top manhattan plots were generated with the variants significant in meta-analysis but not in Korean population nor Japanese population (at p-value < 5 x 10^-8^), meta-analysis using METAL, GENIE and Japanese in the order from top to bottom. The top phenotypes that are uniquely significant by meta-analysis were aPTT, MCHC, DM_diagnosis, K and URIC_ACID.
